# CausalGRN: deciphering causal gene regulatory networks from single-cell CRISPR screens

**DOI:** 10.64898/2025.12.30.692369

**Authors:** Bo Yu, Dingyu Liu, Guanghao Qi, Danwei Huangfu, Li Hsu, Ali Shojaie, Wei Sun

**Author notes:** Contributing authors.

## Abstract

Large-scale single-cell CRISPR screens with single-cell RNA-seq (scRNA-seq) readouts provide critical data to map causal gene regulatory networks (GRNs). However, translating the complex scRNA-seq outputs into reliable causal insights remains a major analytical challenge. Here we present CausalGRN, a scalable computational framework that infers causal GRNs and predicts cellular responses to unseen perturbations. CausalGRN first mitigates pervasive spurious partial correlations in sparse scRNA-seq data through a novel adaptive thresholding correction, enabling robust inference of an undirected graph. It then orients this graph using observed perturbation outcomes. The resulting directed GRN can be used to predict the downstream effects of novel perturbations via network propagation. Across both simulations and diverse experimental datasets, CausalGRN substantially outperforms existing approaches in network reconstruction accuracy and in predicting the effects of unseen perturbations, providing a principled bridge from perturbation data to causal gene regulation.

## Introduction

Understanding the gene regulatory networks (GRNs) that orchestrate cellular function is a central goal of modern biology. A complete and causal map of this circuitry would yield deep insights into development and disease, establishing a predictive model for therapeutic discovery. The emergence of large-scale single-cell CRISPR screens, which systematically perturb genes and measure their transcriptomic consequences using single-cell RNA-seq (scRNA-seq) data, offers an unprecedented opportunity to achieve this vision ^1,2^.

Existing computational frameworks cannot infer GRNs while fully leveraging the scRNA-seq data generated from CRISPR screens. Current approaches fall into two broad categories. The first relies on correlation-based methods, such as tree-based association analyses used in SCENIC ^3–5^. These approaches are not designed to infer causality from perturbation data; instead, they assume a transcription factor regulates its target genes, and infer such relationships solely by association signals. The second category adopts classical causal inference frameworks ^6–8^, which in principle can uncover causal relationships. However, these methods have poor scalability to genome-scale networks. Furthermore, they either rely on restrictive statistical assumptions, such as Gaussianity, that are violated in sparse scRNA-seq count data, or require complicated tests ^9–12^ with low power and poor finite sample performance. These limitations also plague methods for inferring GRNs from low-throughput genetic perturbations using bulk gene expression measurements ^13–15^.

A few recent studies have uncovered a surprising observation: perturbations of thousands of genes can produce transcriptional effects that are similar to one another. Consequently, the average effect across many training perturbations can often give a reasonable prediction of the perturbation effect from unseen perturbations ^16,17^. This observation suggests that distinct perturbations may elicit shared transcriptional programs influencing large portions of the transcriptome. Accounting for such shared effects is therefore essential for accurate GRN inference.

To address these challenges, we present CausalGRN, a scalable computational framework that infers causal GRNs from single-cell CRISPR screens. CausalGRN has four key innovations. First, it employs a novel strategy to infer a network skeleton while minimizing false positives arising from the sparsity of scRNA-seq data. Second, it uses perturbation effects to infer causal relations. Third, it incorporates a gene program within the GRN to model shared perturbation effects. Fourth, CausalGRN enables accurate prediction of perturbation responses by propagating the perturbation effects through the inferred GRN. In addition to CausalGRN, we have also developed GRN-scPerturbSim, a realistic simulator for single-cell perturbation data with a known ground-truth GRN.

Across simulations using GRN-scPerturbSim and diverse experimental datasets, CausalGRN consistently outperforms existing methods in both network inference and perturbation effect prediction. Together, CausalGRN and GRN-scPerturbSim form a comprehensive, validated toolkit that bridges complex perturbation data to causal insight and predictive modeling.

## Results

### Overview

CausalGRN infers causal GRNs through a two-stage process. It first constructs an undirected network skeleton, then orients its edges using the observed outcomes of experimental perturbations (Fig. 1a). The first stage builds the network skeleton using a scalable variant of the Peter-Clark (PC) algorithm, which is based on the principle that conditional independence between two genes implies the absence of a direct causal link between them ^6,18^. The standard PC algorithm assesses conditional independence by partial correlation tests, which fail in scRNA-seq data and lead to widespread spurious partial correlations. This is because the measurement error inherent in the count nature of sparse scRNA-seq data ^19^ leads to weak correlations between underlying and observed expressions for lowly expressed genes, thereby making it very challenging to model the conditional distribution of gene expression (Fig. 1d,e). To mitigate this problem, we introduce a novel method: Adaptive Thresholding Correction for Partial Correlation (ATC-PC), which computes partial correlations only within the subset of cells where the conditioning gene is reliably measured (Fig. 1f; see Methods). Furthermore, to achieve scalability, CausalGRN constrains the conditional-independence search space, an assumption justified by the intrinsic sparsity of GRNs ^20–22^ (see Methods). This solution improves computational efficiency and enables CausalGRN to infer the skeleton from the full aggregate of wild-type and perturbed cells at modern atlas scales, thereby maximizing statistical power.

**Fig. 1.**
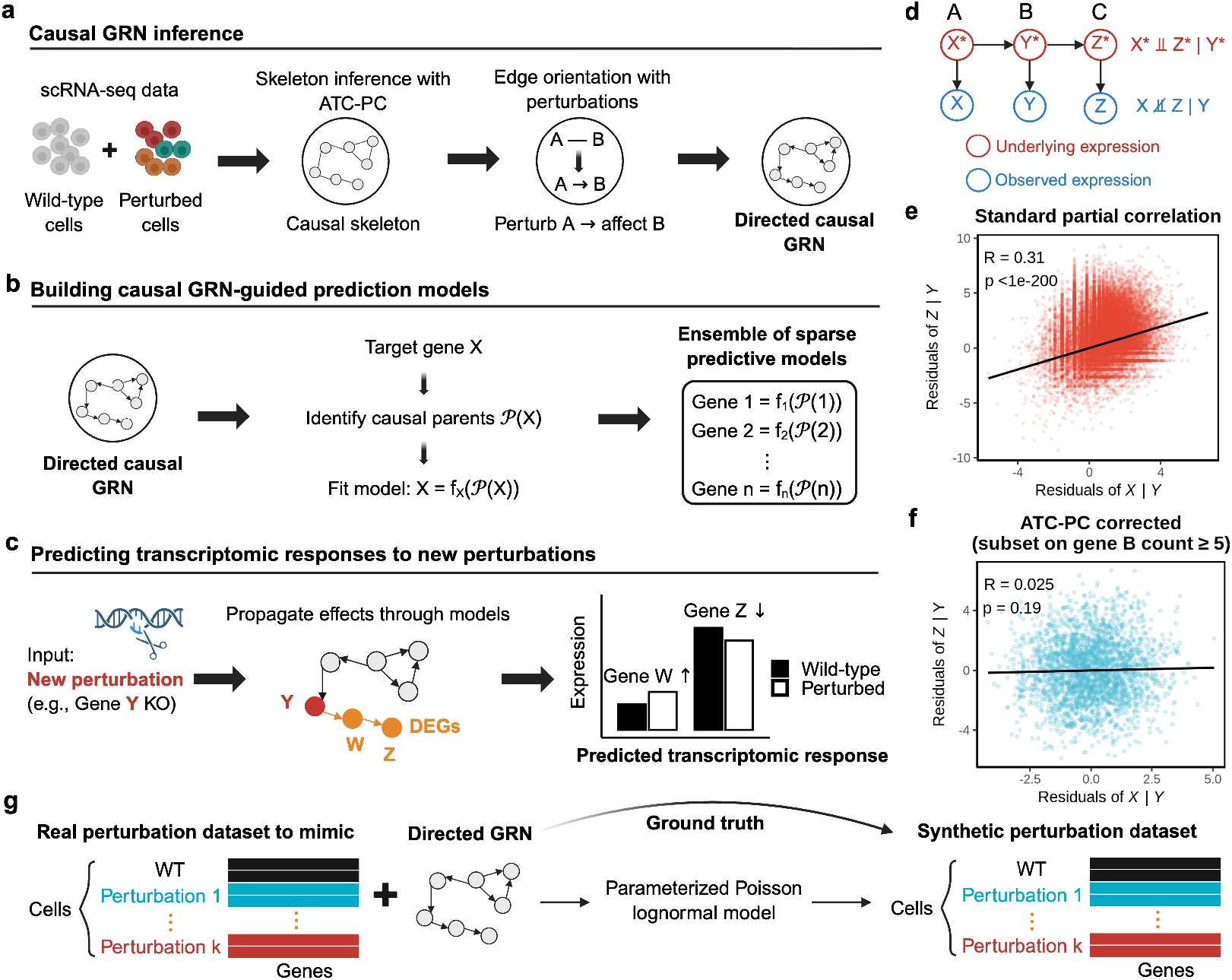
The CausalGRN framework for causal inference of GRNs and perturbation effect prediction. **a**, The causal inference workflow, where a causal skeleton is learned using Adaptive Thresholding Correction for Partial Correlation (ATC-PC) and subsequently oriented using observed perturbation outcomes. An undirected edge *A* − *B* is oriented as *A* → *B* if perturbing A significantly changes B; otherwise it remains undirected. When both genes are perturbed, we assign *A* → *B* only if perturbing A significantly changes B but perturbing B does not significantly change A. **b**, Construction of gene expression prediction models guided by the inferred causal GRN. **c**, Prediction of transcriptomic responses by propagating the effects of new perturbations through the learned prediction models. **d-f**, Correcting spurious partial correlations from measurement error. For a mediated pathway where gene A regulates B and B regulates C, we denote their true underlying expression as *X*^∗^, *Y* ^∗^, *Z*^∗^ and their observed expression as *X, Y, Z* (**d**). *X* and *Z* are independent given *Y* ^∗^, but they are not independent given *Y*, due to measurement error in the observed mediator *Y* (**e**). This artifact is corrected by ATC-PC (**f** ). **g**, The GRN-scPerturbSim workflow, which generates a synthetic perturbation dataset by mimicking a real-world perturbation dataset while enforcing a known, user-provided GRN as the ground truth.

In the second stage, CausalGRN orients the edges of the skeleton by exploiting direct causal evidence from perturbation experiments. The orientation rule is straightforward: an undirected edge *A* − *B* is directed as *A* → *B* if perturbing gene A induces a significant expression change in B (Fig. 1a). This rule is applied systematically across all perturbations, and the resulting high-confidence directions are propagated to orient neighboring edges (see Methods). This principled integration of perturbation data yields a partially directed and causally informed GRN.

Rigorous assessment of CausalGRN’s performance requires realistic single-cell perturbation data paired with a known ground-truth GRN. Since existing simulators are insufficient for this specific task, we have developed GRN-scPerturbSim, a simulator specifically designed for benchmarking causal GRN inference (Extended Data Table 1; see Methods). GRN-scPerturbSim generates realistic single-cell perturbation data guided by a multi-layer ground-truth GRN (Fig. 1g). It reproduces key statistical characteristics of real single-cell perturbation datasets, including sample sizes, gene-specific sparsity, gene-gene correlation structure, and perturbation effect patterns (Extended Data Fig. 1; see Methods).

Finally, CausalGRN supports a predictive modeling framework that uses the inferred causal GRN to predict transcriptional responses to unseen perturbations. For each gene, its expression is modeled as a function of its inferred regulators (Fig. 1b). Given a new perturbation, the initial expression change is propagated through the network to predict the global transcriptomic response (Fig. 1c; see Methods).

### Adaptive thresholding corrects widespread spurious partial correlations in scRNA-seq data

We first used a simulation to quantitatively characterize the severity of the spurious partial correlation artifact and to validate the performance of ATC-PC. We generated datasets from two causal topologies, *A* → *B* → *C* and *A* ← *B* → *C*, where A and C are conditionally independent given B (see Methods). We varied the sparsity level of gene B (Extended Data Fig. 2a) to assess its impact on the Type I error rate for the A–C pair and the statistical power for the A–B and B–C pairs.

The simulation revealed that standard partial correlation testing methods have severe inflation of Type I error when assessing conditional independence between genes A and C given gene B (Fig. 2a). At a moderate sparsity level for gene B, where ∼ 41% of cells had zero counts, standard methods produced type I error rates exceeding 60%, which further escalated to 88% under higher sparsity ( ∼ 92% zero counts). This led to a correspondingly high False Discovery Rate (FDR) (Fig. 2b). In contrast, ATC-PC maintained both Type I error and FDR controls. This rigorous error control was achieved with only a minor reduction in statistical power for detecting the true edges A–B and B–C (Fig. 2c,d). Similar results were observed when using non-parametric Spearman correlation (Extended Data Fig. 2b).

**Fig. 2.**
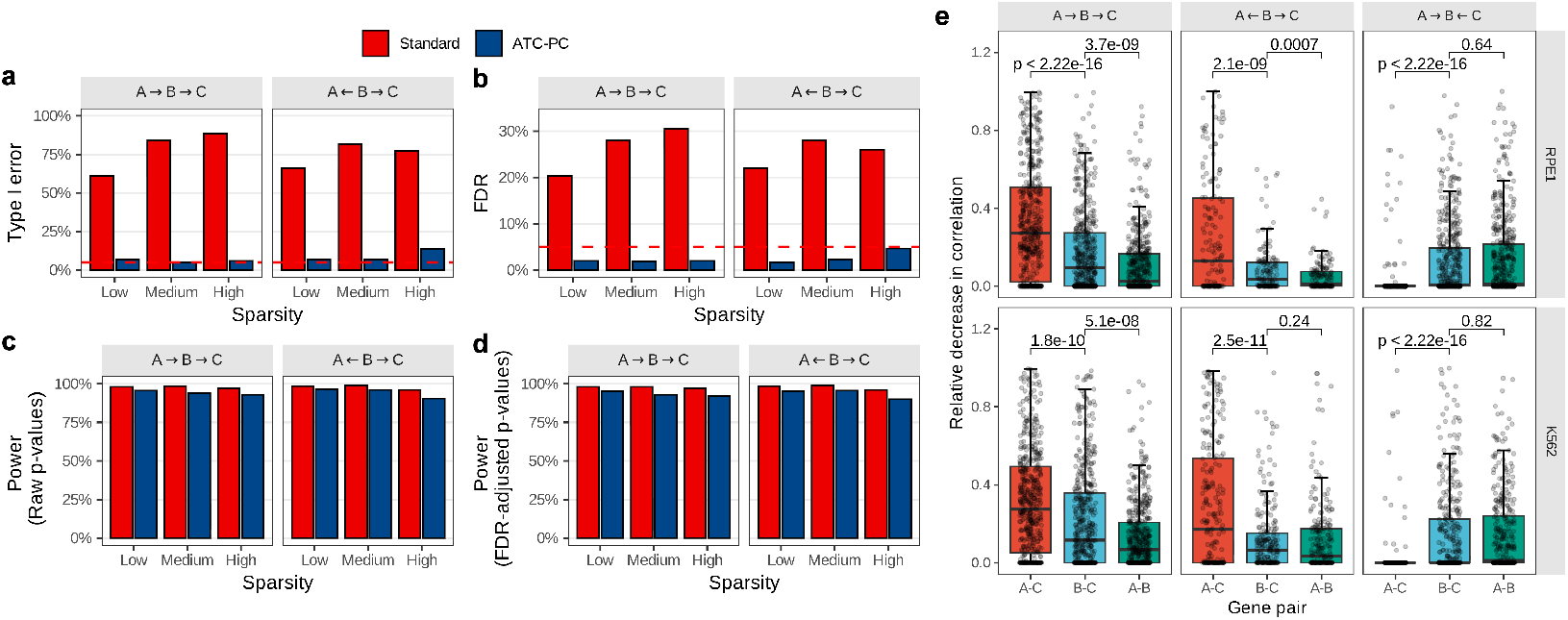
Adaptive thresholding corrects widespread spurious partial correlations in GRN inference from scRNA-seq data. **a-d**, Performance of ATC-PC compared to standard partial correlation in simulated data. Data were generated from two causal motifs (*A* → *B* → *C* and *A* ← *B* → *C*), where A and C are conditionally independent given B. The sparsity of the intermediate gene B was varied from low ( ∼ 41% zero counts) to high ( ∼ 92% zero counts). **a**, Type I error rate for detecting the spurious A–C edge. The dashed red line indicates the nominal 0.05 significance level. **b**, Corresponding FDR for the A–C edge. **c**,**d**, Statistical power to detect the true A–B and B–C edges using raw (**c**) and FDR-adjusted (**d**) *p*-values. All results were generated using Pearson correlation on log-transformed counts, and all metrics were calculated from 100 simulation replicates. **e**, Performance of ATC-PC on gold-standard causal motifs constructed from the RPE1 (top) and K562 (bottom) perturbation datasets. Boxplots show the relative decrease in correlation for the non-adjacent pair (A–C) versus the true edges (A–B and B–C) within chain (*A* → *B* → *C*), fork (*A* ← *B* → *C*), and collider (*A* → *B* ← *C*) motifs. The results align with causal theory: the A–C correlation is significantly attenuated when conditioning on B for chains and forks, while for colliders, it is centered at zero. *P* -values were calculated from two-sided, paired Wilcoxon signed-rank tests.

Next, to evaluate ATC-PC on real data, we developed a validation strategy based on gold-standard causal motifs derived from high-confidence perturbation evidence in the RPE1 and K562 datasets ^23,24^. The key idea was to use perturbation effects as causal anchors. For example, a chain motif (*A* → *B* → *C*) was defined from high-confidence cases where perturbing A significantly affected B’s expression, and perturbing B in turn affected C’s expression. This strategy circumvents the lack of a comprehensive ground-truth GRN by directly testing whether ATC-PC’s statistical behavior on known causal motifs aligns with theoretical expectations. We constructed motifs for the three canonical causal topologies: chains (*A* → *B* → *C*), forks (*A* ← *B* → *C*), and colliders (*A* → *B* ← *C*) (see Methods for details).

Causal theory dictates distinct statistical signatures for these topologies: the A–C correlation should be substantially attenuated by conditioning on gene B in chains and forks, but not in colliders. We therefore applied ATC-PC to these motifs and measured this attenuation by quantifying a relative decrease score in correlation strength (see Methods for formal definition). The results from ATC-PC closely matched the theoretical expectations (Fig. 2e). For chains and forks, conditioning on the intermediate gene B resulted in a large attenuation of the A–C correlation. In contrast, the relative decreases for the true edges B–C (i.e., B–C given A versus B–C) and A–B (i.e., A–B given C versus A–B) were substantially smaller. Conversely, for colliders, where A and C were independent causes of B, most of the relative decreases for A–C were close to zero, consistent with the expectation since A–C became dependent given B and the score correctly registered no attenuation. These results provide strong evidence for ATC-PC’s ability to reliably resolve local causal structures within complex GRNs.

### CausalGRN substantially improves GRN reconstruction in simulations

CausalGRN overcomes the fundamental limitations of existing methods for causal GRN inference from single-cell perturbation data (Extended Data Table 2). We first demonstrate this with a simulated three-gene chain (Fig. 3a), where CausalGRN correctly identifies the ground-truth network (Fig. 3b) while the two major classes of existing methods fail as expected (Fig. 3c,d). Specifically, association-based methods fail to distinguish causality from mere correlation, while classical causal inference methods are unsuitable due to the inherent sparsity of scRNA-seq data (see Extended Data Fig. 3 for details).

**Fig. 3.**
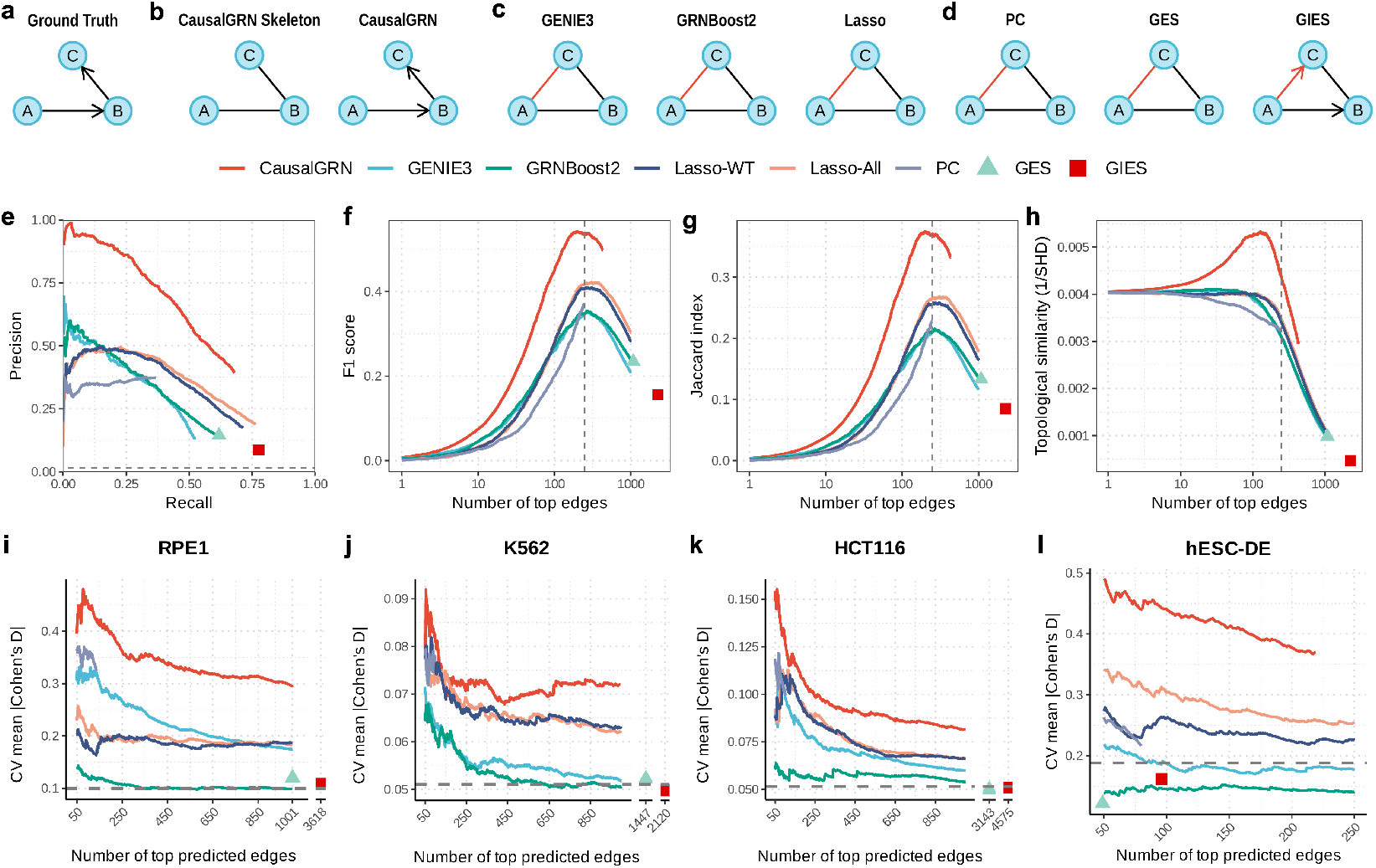
CausalGRN outperforms existing methods on both simulated and experimental data. **a–d**, Evaluation on a simulated three-gene chain (*A* → *B* → *C*) demonstrating the failure of existing methods. Edges matching the ground truth are colored black, and incorrect edges are red. See Extended Data Fig. 3 for details. **a**, The ground-truth network. **b**, CausalGRN correctly infers the skeleton and orients all edges using the perturbation data. **c**, Association-based methods incorrectly infer a direct link between A and C, failing to distinguish causality from correlation. **d**, Classical causal inference methods fail to recover the network, as their models are unsuitable for sparse scRNA-seq data. **e–h**, Performance on the GRN-scPerturbSim benchmark, averaged across ten replicates. Lasso-WT: Lasso applied to wild-type cells; Lasso-All: Lasso applied to all cells (wild-type and perturbed). **e**, Precision-Recall curve; the horizontal dashed line indicates random precision. **f**, F1 score. **g**, Jaccard index. **h**, Topological similarity (1/SHD), where SHD is Structural Hamming Distance. In **f–h**, metrics are plotted versus the number of top-ranked edges, and the vertical dashed line indicates the average number of true edges in the ground-truth networks. Methods returning a single unweighted graph (GES, GIES) are plotted as single points. **i–l**, Performance on four experimental datasets, evaluated using a five-fold cross-validation. The y-axis, “CV mean |Cohen’s D |”, measures the mean causal evidence for the top *N* predicted edges (x-axis). The datasets are RPE1 (**i**), K562 (**j**), HCT116 (**k**) and hESC-DE (**l**). Methods returning a single unweighted graph (GES, GIES) are plotted as single points.

To confirm these conceptual advantages at scale, we benchmarked CausalGRN against seven representative methods using our GRN-scPerturbSim simulator, which generated a realistic testbed recapitulating the key statistical properties of the RPE1 dataset (Extended Data Fig. 1; see Methods).

CausalGRN substantially outperformed all seven competitors in reconstructing the ground-truth networks. Averaged across ten replicate simulations, CausalGRN was the top performer across all four evaluation metrics, which assessed both the overall quality of the ranked edge list and the accuracy at various cutoffs (Fig. 3e–h; see Methods). These results provide direct evidence that CausalGRN’s principled design is highly effective. The superior performance is not simply due to the use of interventional data, as CausalGRN also substantially outperformed other methods that also leverage the perturbed cells, such as GIES and Lasso-All. An ablation study further confirmed that each component of CausalGRN contributes synergistically to the final accuracy (Extended Data Fig. 4).

### CausalGRN identifies causal links from large-scale experimental data

We next benchmarked CausalGRN using four experimental datasets from diverse cellular contexts: three large-scale Perturb-seq screens in human RPE1, K562^23,24^, and HCT116^25,26^ cell lines, and a smaller CRISPR knockout screen in definitive endoderm differentiated from human embryonic stem cells ^27^ (hESC-DE; see Methods). To ensure rigorous benchmarking, we assembled a high-quality perturbation set for each dataset by filtering for perturbations with sufficient cells to reliably measure perturbation effects, totaling 639 perturbations (see Methods).

Accurate GRN inference in these datasets requires explicitly modeling gene programs as latent factors that represent specific cellular activities and thereby capture shared perturbation effects on their downstream genes. Ignoring these gene programs leads to spurious edges between genes co-regulated by gene programs, while regressing out gene program signatures induces artificial correlations among their upstream regulators (Extended Data Fig. 5a, b). CausalGRN overcomes this challenge by explicitly incorporating gene programs into the network structure. We first empirically characterized these programs by analyzing coordinated transcriptional responses to perturbations, confirming that the dominant response program is robustly captured by a proxy: the first principal component of wild-type gene expression (WT-PC1; see Methods). CausalGRN therefore incorporates WT-PC1 as a “pseudo-gene” node into the GRN.

We benchmarked all eight methods using a five-fold cross-validation scheme to assess the causal validity of inferred edges. This evaluation is based on the principle that a predicted edge *A* → *B* should correspond to a measurable change in B when A is perturbed. In each fold, we inferred the network from the training data and assessed the subset of inferred edges whose source gene corresponded to a held-out perturbation. The experimental evidence supporting each edge was quantified using the absolute value of Cohen’s D, comparing the target gene’s expression between wildtype cells and cells in which the source gene was perturbed. For consistency across methods, the CausalGRN network, which internally models the WT-PC1 pseudo-gene, was projected back into the gene-only space for evaluation (see Methods).

The experimental results corroborated our simulation findings (Fig. 3i–l). CausalGRN consistently outperformed all seven existing methods across all four datasets. The cross-validated mean |Cohen’s D| for CausalGRN was substantially higher at all evaluated cutoffs, indicating that its top-ranked edges correspond to stronger and more reproducible causal effects. These results demonstrate that CausalGRN’s principled framework yields substantially more accurate causal GRNs from complex single-cell perturbation data.

### CausalGRN enables accurate perturbation effect prediction in simulations

We evaluated the CausalGRN-guided predictive framework using the GRN-scPerturbSim simulated dataset under a five-fold cross-validation scheme (see Methods). Two strong baselines were included for comparison. The first baseline, Average Effect, predicts each test perturbation’s outcome as the mean perturbation effect observed in the training set, a simple yet surprisingly competitive approach that was recently shown to outperform several large foundation models ^16,17^. The second baseline, Complete Graph-guided prediction, uses the same propagation algorithm as the CausalGRN-guided framework but fits per-gene expression models via Lasso on all other genes. Therefore, in contrast to Complete Graph-guided predictions, CausalGRN-guided framework exploits the value of the inferred GRN.

CausalGRN-guided prediction consistently outperformed both baselines, yielding higher precision for identifying up- and down-regulated DEGs across all thresholds (Fig. 4a). This advantage comes from its causal design: it propagates perturbation effects strictly along inferred “parent → child” directions in the network. By contrast, the Complete Graph-guided framework diffuses signals across an association network, which invites non-causal errors such as reverse causation (i.e., propagating from a child to its parents), while the Average Effect baseline cannot capture perturbation-specific effects by design.

**Fig. 4.**
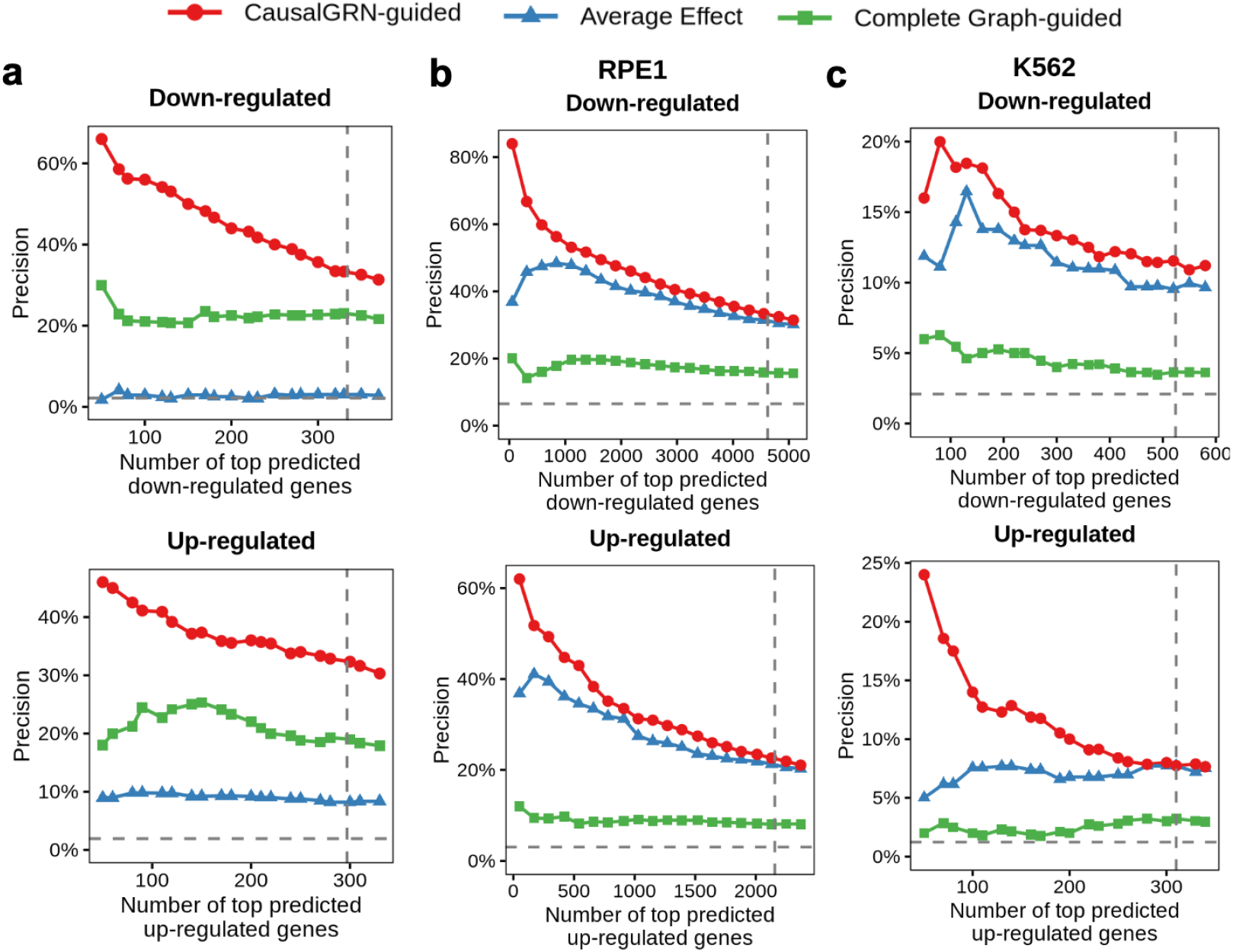
CausalGRN enables superior prediction of DEGs from unseen perturbations. Precision in identifying true down-regulated (top row) and up-regulated (bottom row) DEGs from held-out perturbations in five-fold cross-validation. Predictions from all folds were pooled to generate each curve. Panels show results for (**a**) the GRN-scPerturbSim simulated dataset, (**b**) RPE1, and (**c**) K562. Precision (y-axis) is the fraction of true down- (or up-) regulated DEGs found within the top *N* genes (x-axis) ranked by the most negative (or positive) predicted differential expression, respectively. The plot compares three prediction frameworks: CausalGRN-guided, which builds expression models using linear regression on inferred causal parents and applies the perturbation effect propagation algorithm (see Methods); the Average Effect baseline, which predicts the training-set average perturbation effect for all test perturbations; and Complete Graph-guided, which builds expression models using Lasso regression on all other genes and applies the same propagation algorithm. The horizontal dashed line indicates random precision; the vertical dashed line marks the total number of true down- or up-regulated DEGs in the pooled held-out test set.

Both the Complete Graph-guided and CausalGRN-guided frameworks predict perturbation effects on each gene using two components: (i) a per-gene expression model and (ii) estimated values of its predictor genes. Performance thus depends on the accuracy of both components. To dissect why CausalGRN-guided prediction outperformed the Complete Graph-guided baseline, we evaluated both in an oracle setting, where predictor gene expression was directly observed from the test perturbation and used to forecast the target. In this ideal scenario, Lasso models of the Complete Graph-guided framework achieved similar or higher accuracy than CausalGRN-guided models (Extended Data Fig. 6a). This result contrasts sharply with its poor performance in the practical setting, where predictor values are unknown and must be inferred by propagating perturbation effects. This gap demonstrates that propagation fails without the causal scaffold, regardless of the complexity of gene expression prediction models.

### CausalGRN enables accurate perturbation effect prediction in real-world datasets

We next evaluated the CausalGRN-guided predictive framework on the large-scale RPE1, K562 and HCT116 datasets. The hESC-DE dataset was excluded because its 14 perturbations were insufficient to infer a GRN with enough directed edges for effective propagation of perturbation effects. CausalGRN-guided prediction was benchmarked against the Complete Graph-guided prediction and the Average Effect baseline. Consistent with previous reports ^16,17^, the Average Effect baseline performed well in these datasets. Our analysis provided an explanation for this observation: many perturbations induced similar responses in the WT-PC1 program, which in turn drove a coordinated downstream transcriptional response (Extended Data Fig. 7a).

For CausalGRN-guided prediction, we used the inferred GRN that included the WT-PC1 program as a node. The response of the WT-PC1 program represents a complex, system-wide effect which cannot be modeled by simple gene-gene regulations. To model this component appropriately, we employed a specialized portability model for this single node, leveraging our observation that the program’s response was a transferable feature across datasets (Extended Data Fig. 7b–d). The predicted response of the WT-PC1 program was then combined with the direct effect on the perturbed gene to define the initial signal, which was subsequently propagated through the causal scaffold to predict the expression changes of all downstream genes.

This integrated framework, which combines gene program information with a causal scaffold for propagation, proved highly effective. In the RPE1 and K562 datasets, CausalGRN-guided prediction showed a clear and consistent advantage, outperforming both baselines in predicting up- and down-regulated DEGs (Fig. 4b,c). In contrast, the Average Effect baseline fails to capture perturbation-specific responses, and the Complete Graph-guided baseline cannot propagate effects along causal cascades.

In the HCT116 dataset, however, no method was consistently superior (Extended Data Fig. 7e). This observation aligned with the finding that the dominant gene program in HCT116 was less well represented by its WT-PC1 proxy (Extended Data Fig. 5j), and that the WT-PC1 response exhibited lower portability from the reference datasets (Extended Data Fig. 7d), thereby diminishing the advantage of our framework.

Across the RPE1, K562 and HCT116 datasets, the oracle analysis reproduced our simulation findings for perturbation effect prediction: the Complete Graph–guided framework achieved high accuracy when true predictor values from test perturbations were supplied (Extended Data Fig. 6b–d), indicating that its Lasso-based expression models were sufficient. Its failure in the practical setting again highlighted the catastrophic failure of propagation on a non-causal graph. Overall, the results show that accurate perturbation effect prediction in real data requires both a causal scaffold for correct propagation and the explicit, careful modeling of gene programs.

### CausalGRN resolves regulatory hierarchies and prioritizes upstream regulators of Type 2 diabetes risk genes

To dissect the upstream regulation of Type 2 Diabetes (T2D) risk genes, we applied CausalGRN to a single-cell CRISPR screen in hESC-derived beta cells ^27^, a human differentiation model for interrogating beta-cell regulatory programs relevant to insulin secretion and T2D (see Methods). We modeled a network of 327 genes, defined as the union of 312 T2D risk genes, 8 knockout targets, and 8 additional developmental genes.

First, we benchmarked the inferred topology against a correlation network generated by WGCNA ^28^. At matched edge density, the correlation network captures undirected, dense co-expression links concentrated around program genes, whereas CausalGRN resolves partially directed hierarchies around knockout genes (Fig. 5a). We assessed the accuracy of these inferred interactions against the DoRothEA database ^29^. Despite limited ground-truth coverage for this specialized gene set, 8.1% of the CausalGRN edges overlapped with validated interactions from DoRothEA, whereas the correlation network recovered zero validated interactions. For *FOXA1*, the most well-annotated transcription factor by DoRothEA in the network, CausalGRN correctly recovered three annotated targets out of three predicted children (100% precision), compared to none for WGCNA. For instance, the topology places the pioneer factor *FOXA1* upstream of the metabolic regulator *LDHB*. This ordering aligns with the established role of FOXA factors in maintaining beta-cell metabolic competency, identifying the metabolic effector *LDHB* as a candidate downstream node consistent with this regulatory program ^30,31^.

**Fig. 5.**
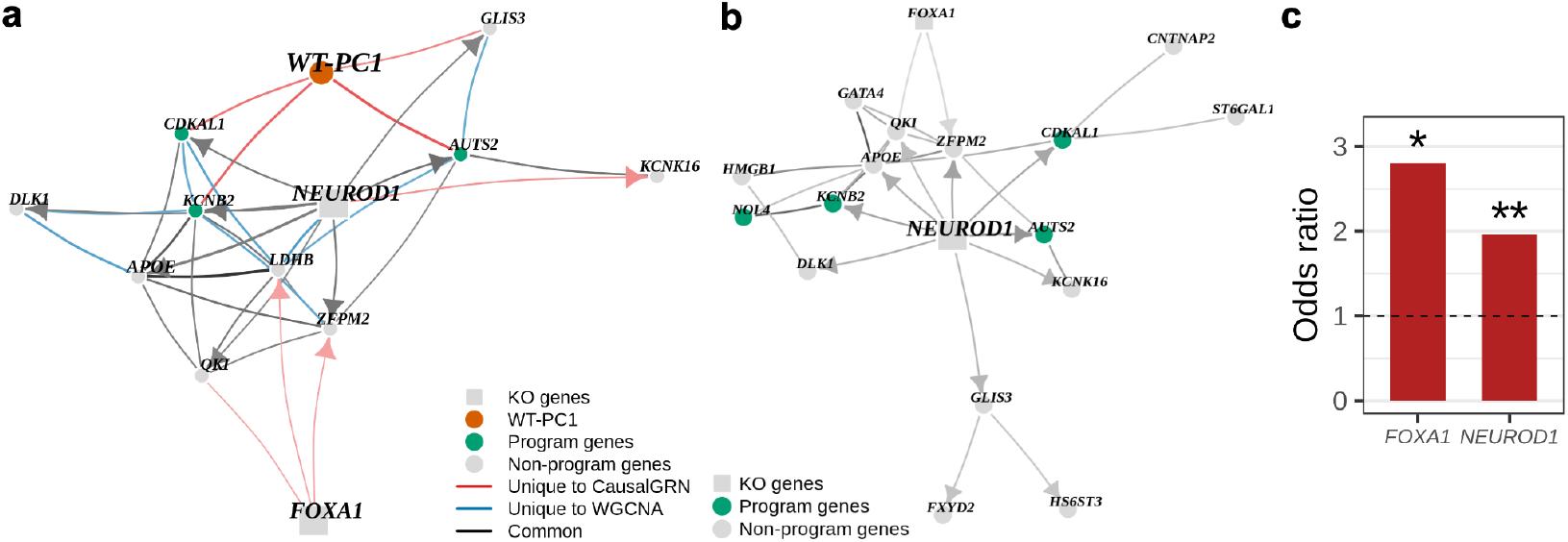
CausalGRN resolves regulatory hierarchies and prioritizes upstream regulators of T2D risk genes. **a**, Comparison of CausalGRN and WGCNA edges on a subnetwork around knockout genes in hESC-derived beta cells. At matched edge density, WGCNA captures undirected co-expression relationships concentrated around program genes, whereas CausalGRN resolves partially directed hierarchies around knockout genes. Edge width and transparency reflect edge strength (CausalGRN confidence score for CausalGRN-only and common edges (see Methods); co-expression strength for WGCNA-only edges). **b**, CausalGRN converts *NEUROD1* -knockout DEGs from a flat list into a structured architecture. The subnetwork visualizes *NEUROD1* -knockout DEGs, restricted to the union of the top 15 DEGs and all CausalGRN-inferred children of *NEUROD1*. **c**, CausalGRN prioritizes upstream regulators of T2D risk genes by descendant enrichment. Each bar shows the enrichment of T2D risk genes (relative to mean-expression-matched control genes) among the descendants of each knockout gene in the CausalGRN network (see Methods). Only *FOXA1* and *NEUROD1* show significant enrichment after Benjamini-Hochberg correction (∗: adjusted *p*-value *<* 0.05, ∗∗: adjusted *p*-value *<* 0.01).

Second, the inferred topology enables a more mechanistic dissection of perturbation effects. It converts *NEUROD1* -knockout DEGs from a flat list into a causal hierarchy (Fig. 5b). For example, it resolves a multi-step regulatory path: *NEUROD*1 → *GLIS*3 → *FXY D*2. The placement of *GLIS3* downstream of *NEUROD1* is consistent with their participation in shared islet regulatory hubs, where *GLIS3* co-occupies regulatory loci with *NEUROD1* and can functionally cooperate with *NEUROD1* to activate core beta-cell transcriptional programs ^32,33^. Extending this axis, the inferred topology predicts *FXYD2* as a downstream target of *GLIS3*, nominating a specific regulatory link between *GLIS3*, a maturation-relevant islet regulator ^32^, and *FXYD2*, a maturation-associated marker and regulator ^34^. Thus, the inferred topology resolves these gene-level effects into a coherent maturity-associated module, providing concrete mechanistic hypotheses for experimental validation.

Finally, we prioritized upstream regulators of the T2D risk genes using a descendant enrichment analysis. To establish a statistical background for the enrichment tests, we expanded the network to include non-risk control genes selected to match the average expression of the risk genes. We then tested each knockout gene for the enrichment of T2D risk genes among its descendants relative to these controls (see Methods). This topology-driven analysis identified *FOXA1* and *NEUROD1* as significant drivers after Benjamini-Hochberg correction (Fig. 5c). Together, these results demonstrate how CausalGRN distills high-dimensional perturbation data into interpretable causal topologies, enabling systematic prioritization of upstream drivers in disease-relevant gene networks.

## Discussion

We developed CausalGRN, a scalable, principled framework for inferring causal GRNs from large-scale single-cell CRISPR screens. This powerful data modality poses two central methodological challenges: (i) scalable inference of causal network structure from sparse, high-dimensional data, and (ii) accurate prediction of cellular responses to new perturbations. CausalGRN addresses both challenges through a multi-stage causal inference framework. In parallel, we provide GRN-scPerturbSim as a valuable community resource for benchmarking future GRN inference methods.

CausalGRN’s foundation is a scalable variant of the PC algorithm that infers network structure via conditional independence testing. To adapt this classical approach to sparse scRNA-seq data, CausalGRN introduces ATC-PC, which corrects widespread spurious partial correlations and enables robust reconstruction of the network skeleton. In genome-wide perturbation studies, where distinct gene perturbations often trigger shared transcriptional programs, CausalGRN models the dominant shared response as a “pseudo-gene” node. This step effectively removes thousands of false edges among co-regulated genes. Finally, CausalGRN orients the resulting skeleton using interventional data. This multi-pronged strategy, combining artifact correction (ATC-PC), gene program modeling, and perturbation-guided orientation, enables CausalGRN to build a reliable causal scaffold from CRISPR screens.

Beyond GRN inference, the CausalGRN-guided predictive framework offers a new perspective on the challenge of predicting perturbation effects. The unexpected strength of a simple Average Effect baseline, which outperformed large foundation models in predicting perturbation effects, has remained puzzling ^16,17^. Our results provide a mechanistic explanation: this Average Effect baseline performs well because many perturbations elicit similar changes in a dominant gene program. The challenge is to identify such a gene program before perturbing thousands of genes. We found that the first principal component from wild-type data (WT-PC1) can capture this dominant gene program. The CausalGRN-guided prediction framework explicitly models this gene program, leveraging our finding that this program is a portable feature across datasets. The importance of the causal scaffold from CausalGRN is underscored by the failure of the Complete Graph-guided method, which uses as many genes as possible for prediction, regardless of their regulatory relationships. In contrast, CausalGRN only uses parent nodes to predict their child nodes.

Several avenues exist to improve CausalGRN. First, its performance is linked to a user-provided representation of gene programs, which requires manual examination. In our studies, we represented the dominant program with WT-PC1, which was a good proxy in the RPE1, K562, and hESC-DE datasets, but it had a weaker association with the dominant program in the HCT116 dataset. Second, the CausalGRN-guided predictive framework requires suitable reference data from cell systems where the target dataset’s program is also present. Third, the inferred network is only partially directed, as orientation depends entirely on the set of available perturbations. Finally, CausalGRN’s skeleton inference constrains the search space to achieve scalability, which may miss higher-order conditional independencies.

These limitations point to clear future directions. Methodologically, CausalGRN could be extended to automatically identify and represent essential gene programs, removing the dependency on manual characterization. The prediction framework’s data dependence highlights a second key opportunity: a systematic, large-scale investigation of gene program response portability. By leveraging growing public perturbation databases ^35,36^, one could build a “Portability Atlas” to quantify the similarity of gene program responses to a perturbation across hundreds of cellular contexts. A separate, and perhaps larger, opportunity lies in tackling the portability of the GRN structure itself: transferring a learned network from a data-rich system to a new context. To address the incomplete orientation, CausalGRN could also integrate complementary data types, such as chromatin accessibility (ATAC-seq) or expression quantitative trait loci (eQTLs), as orthogonal evidence. Finally, from an application perspective, CausalGRN can rank genes by how often they are predicted to lie downstream of genes whose perturbations produce similar cellular phenotypic consequences. Genes that repeatedly emerge as shared downstream targets may sit closer to the mechanisms underlying the shared consequence and thus provide actionable candidates for follow-up validation.

CausalGRN provides a robust, scalable suite of tools that moves the field beyond simple correlation networks or black-box prediction. By tackling the dual challenges of inference and prediction within a single, unified causal framework, it offers a principled path from complex single-cell perturbation data to the two central goals of systems biology: deep causal insight and true predictive power. The CausalGRN framework is poised to tackle a wide range of questions, such as deciphering the networks driving cell differentiation, identifying mechanisms of drug resistance, or mapping the causal circuitry of genetic diseases.

## Methods

### Single-cell perturbation datasets

#### RPE1 and K562 datasets

We used two large-scale Perturb-seq datasets derived from human RPE1 and K562 cell lines ^23^. The raw counts, pre-normalized expression z-scores, and associated metadata were downloaded from Figshare ^24^ and the original publication’s supplementary information.

For both datasets, we retained only strong knockdowns, defined as those with at least 50 perturbed cells, a significant on-target effect ( ≥ 30% reduction in target gene expression, as reported in the source publication), and at least ten differentially expressed genes (DEGs), as determined by the Anderson–Darling test in the source publication. For GRN analysis, we excluded mitochondrial and ribosomal protein genes, because they often reflect global cell state or technical variation and can form spurious hubs that distort GRN inference ^37^. To remove extreme outliers, we clipped pre-normalized expression z-scores to a maximum value of 10. We then regressed out covariates including library size, mitochondrial read percentage, and cell cycle scores (S and G2M phases).

The final RPE1 expression data contained 8,644 genes across 166,062 cells, comprising 11,485 wild-type cells and 154,577 perturbed cells from 1,159 strong knockdowns. The final K562 expression data contained 8,071 genes across 580,498 cells, comprising 75,328 wild-type cells and 505,170 perturbed cells from 2,486 strong knockdowns.

#### HCT116 dataset

This is a large-scale Perturb-seq dataset from the human HCT116 cell line ^25^. The raw count matrix and associated metadata were downloaded from Figshare ^26^. First, we calculated log-normalized expression values by dividing the counts in each cell by its library size, multiplying by a scale factor of 10,000, and applying a log(*x* + 1) transformation. Expression values were subsequently clipped at a maximum of 10.

Second, to correct for technical variability between experimental batches, we adapted the strategy from Replogle et al. ^23^, using the wild-type cells within each batch as a reference. For each batch and each gene, we first calculated the mean and standard deviation of gene expression using only wild-type cells. These batch-specific statistics were then used to standardize gene expression of all cells from the corresponding batch by subtracting the mean and dividing by the standard deviation. This transformation removes batch effects by aligning wild-type cells across batches while preserving the biological difference between perturbed and wild-type cells. In addition, we excluded mitochondrial and ribosomal protein genes, and then regressed out covariates including library size, mitochondrial read percentage, and cell cycle scores (S and G2M phases).

Third, to identify DEGs, we compared expression in perturbed cells to wild-type cells using both a two-sample t-test and a Wilcoxon rank-sum test. We defined DEGs as those with Benjamini-Hochberg adjusted *p*-values *<* 0.05 from both tests. This conservative approach ensures robustness against false positives from the t-test due to violations of normality assumptions and against false positives of the Wilcoxon ranksum test due to small systematic normalization shifts. It is used throughout this study unless otherwise specified.

Finally, we retained strong knockdowns with at least 50 perturbed cells, a significant on-target knockdown (the target gene itself being a DEG), and at least ten DEGs. The final HCT116 data contained 12,419 genes across 435,763 cells, comprising 162,842 wild-type cells and 272,921 perturbed cells from 1,462 strong knockdowns.

#### hESC-DE dataset

We also used a dataset from a pooled CRISPR-Cas9 knockout screen in the H1 and HUES8 human embryonic stem cell (hESC) lines during pancreatic differentiation ^27^. As the perturbation method yields non-functional proteins, we first set the target gene’s counts to zero in perturbed cells. Next, we calculated log-normalized expression values by dividing the counts for each cell by its library size, multiplying by the median library size across all cells, and applying a log(*x* + 1) transformation. Expression values were not clipped, as extreme outliers were not observed. Consistent with our other analyses, we excluded mitochondrial and ribosomal protein genes and regressed out covariates including library size, mitochondrial read percentage, and cell cycle scores (S and G2M phases).

To create a homogeneous cohort for analysis, we focused on day 3 definitive endoderm (DE) cells, thereby reducing cellular heterogeneity from the differentiation protocol. From this population, we selected all single-gene, homozygous knockouts with at least 50 perturbed cells. The final hESC-DE expression data contained 9,321 genes across 8,408 cells, comprising 2,729 wild-type and 5,679 perturbed cells from 14 knockouts.

#### hESC-beta dataset

We applied the processing pipeline described above for the hESC-DE dataset to the hESC-derived beta cell population ^27^. The hESC-beta dataset contained 9,630 genes across 5,164 cells, comprising 3,554 wild-type cells and 1,610 perturbed cells from 8 knockouts.

### Adaptive thresholding correction of partial correlation (ATC-PC)

The standard estimate of partial correlation between gene A and C given B is defined as

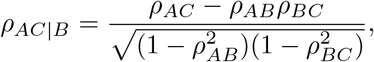

where *ρ*_*AB*_, *ρ*_*AC*_ and *ρ*_*BC*_ are the marginal correlation estimates from the observed expression of genes A, B, and C.

This estimation is problematic for scRNA-seq data, as it implicitly assumes that the observed expression of gene B is a reliable proxy for its latent expression. However, the inherent count nature of scRNA-seq data introduces substantial measurement error ^19^ so that the observed expression is a poor proxy for underlying expression when the underlying expression level is low. Based on the Directed Acyclic Graph (DAG) framework for causal models, A and C are independent given B only if B blocks all paths between A and C. However, the noisy version of B cannot effectively block such paths (Fig. 1d). Consequently, the standard method severely overestimates the partial correlation, leading to inflated type I errors.

To address this issue, ATC-PC implements an adaptive thresholding procedure that selects a subset of cells where the conditioning gene’s expression is high enough to be a reliable proxy for its underlying expression. Specifically, we select cells where the expression of the conditioning gene B exceeds an adaptive threshold *T* . This threshold *T* is the maximum of *t*’s that fulfill two conditions: (i) the expression (i.e., count data) of gene B *> t* in at least *n*_1_ (default 1,000) cells, and (ii) among these cells, at least *n*_2_ (default 200) have nonzero expression for genes A and C. The search for *T* is capped at a maximum value of 10, as counts at this level already provide a reliable proxy for the underlying expression. The second criterion is needed so that this selection does not bias towards the cells where A and C have low expression, particularly in scenarios where the expression of B negatively correlates with that of A and C. Given the selected cells, we perform partial correlation test using Fisher’s *z*-transformation on normalized scRNA-seq data, such as log-transformed counts.

### Causal inference of GRNs with single-cell perturbation data

#### Skeleton inference with ATC-PC

The GRN skeleton is inferred using a parallelized and reduced variant of the PC-Stable algorithm ^18^, which begins with a complete graph and iteratively prunes edges that represent conditional independence. Our implementation is adapted for scRNA-seq data to be both computationally scalable and robust against the statistical artifacts of data sparsity.

To achieve computational scalability, we first constrain the search for conditional independencies to a maximum order of one (i.e., conditioning on only a single gene), a simplification justified by the inherent sparsity of GRNs that makes the process tractable for thousands of genes ^20–22^. Next, by adopting the PC-Stable framework ^18^, all conditional independence tests of a given order are performed in parallel.

Robustness is achieved by employing ATC-PC as the core conditional independence test, which prevents the high rate of false discoveries driven by data sparsity. During the iterative pruning, ATC-PC uses a dual criterion to remove an edge: it is pruned if the partial correlation is not statistically significant or if its magnitude falls below a predefined threshold.

For the edges that remain in the final skeleton, we define a confidence score to enable their ranking. The score for each edge is the minimum absolute partial correlation observed across all conditional independence tests involving that edge. This metric provides a conservative estimate of an edge’s evidence, allowing the final edges to be ranked by confidence. The complete procedure is formally outlined in **Supplementary Algorithm 1**.

#### Causal edge orientation using perturbation data

The skeleton’s undirected edges are oriented using the perturbation data, where the downstream effects of each perturbation provide direct causal evidence. Specifically, an undirected edge A–B is oriented as *A* → *B* if the direct perturbation of gene A causes significant differential expression in B. To resolve ambiguity when both genes in a pair have been perturbed, this orientation is set only if the evidence is one-sided (i.e., perturbing B does not cause a significant change in A). As an optional step to increase the number of directed edges, the algorithm can apply a transitive orientation procedure. This procedure uses the set of edges oriented in the last step to resolve more distant, undirected edges (e.g., orienting *B* → *C* from a path *A* → *B* − *C*; see **Supplementary Note 1** for details). The final output is a partially directed graph representing the inferred causal GRN.

### GRN-guided simulation of single-cell perturbation data (GRN-scPerturbSim)

Rigorous evaluation of GRN inference methods requires realistic single-cell perturbation data coupled with a known ground-truth GRN. However, existing simulators do not fully meet this need (Extended Data Table 1). Data-driven simulators, such as Splatter ^38^ and SPARSim ^39^, accurately reproduce dataset-level statistics but do not embed an explicit underlying GRN. In contrast, trajectory-based simulators, including SERGIO ^40^ and dyngen ^41^, simulate GRN-driven expression dynamics, yet are not directly parameterized to match a specific perturbation screen. Among GRN-driven steady-state simulators, GeneSPIDER2^42^ employs generically parameterized models rather than learning from specific real perturbation screens, while GRouNdGAN ^43^ assumes a bipartite GRN structure, constraining transcription factor interactions and limiting multi-layer regulatory cascades.

To overcome the limitations of existing approaches, we developed GRN-scPerturbSim, a simulator that generates realistic single-cell perturbation data guided by a multi-layer ground-truth GRN and is quantitatively parameterized from real perturbation datasets. GRN-scPerturbSim reproduces key statistical characteristics of the target dataset, including sample sizes, gene-specific sparsity, gene-gene correlation structure, and perturbation effect profiles. This enables a realistic and controlled testbed for benchmarking the accuracy of GRN reconstruction methods.

The simulation proceeds through four stages. First, a ground-truth GRN is generated using the Barabási–Albert model, which yields the scale-free topology characteristic of biological regulatory networks ^44,45^. Second, gene expression models are constructed based on this GRN, with parameters calibrated to reproduce the empirical statistical properties of the target dataset (e.g., gene-specific sparsity, gene–gene correlation distribution, and perturbation effect pattern), ensuring quantitative realism. Third, wild-type cells are generated by sampling from the calibrated model. Finally, perturbations are simulated through a two-stage process: an initial direct effect is applied to the target gene using a tunable perturbation efficacy parameter to capture the incomplete penetrance of CRISPR-based perturbations, followed by propagation of this effect through the network, allowing indirect downstream responses to arise naturally via cascading expression changes. Full mathematical details are provided in **Supplementary Note 2**.

### Predicting transcriptomic responses to unseen perturbations

Our framework predicts the transcriptomic consequences of diverse experimental perturbations, including single- or multi-gene knockouts, knockdowns, and over-expression. To do so, we predict the steady-state transcriptomic response to these *in silico* perturbations using a linear system constrained by an inferred causal GRN. We assume gene expression *X* is governed by *X* = *α* + *BX* + *e*, where *B, α* and *e* are the regulatory coefficient matrix, intercept vector, and noise vector, respectively.

#### Training of expression prediction models

To estimate the parameters *B* and *α*, a separate linear regression model is fit for each gene. The model’s predictors are strictly limited to the gene’s parents in the inferred causal GRN (including neighbors connected by undirected edges). To ensure the model learns each gene’s endogenous regulation, it is trained on all cells except those in which that gene was directly perturbed.

#### Perturbation effect prediction via network propagation

To predict the effect of a new perturbation on a set of genes *I*, we partition the system into three disjoint sets: the perturbed genes *I*, their network descendants *U*, and all other unaltered genes *K*. This partitions the system into a set of unknown expression values to be predicted, *x*_*U*_, and a set of known input values, 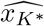 (which combines the specified perturbation values *x*_*I*_ and the wild-type level expression for genes in *K*).

This formulation yields a solvable linear system for the unknown expression values *x*_*U*_ . The unique solution requires the matrix 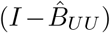 to be invertible and is given by:

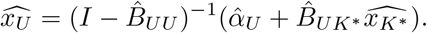

This solution represents---a stable equilibrium if the spectral radius of the internal network, 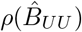, is less than 1. Across all our analyses, this condition was met. This equation propagates the initial, direct effects from the *K*^∗^ genes 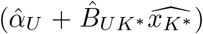 through the internal network of affected genes *U* via the propagator matrix 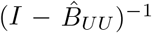.

A complete mathematical derivation, including the partitioning logic and a formal analysis of solution stability, is provided in **Supplementary Note 3**.

### Validating ATC-PC in simulation

This simulation quantifies how standard partial correlation estimation fails on scRNA-seq data and demonstrates the performance of ATC-PC in controlling type I error and FDR while retaining power.

#### Simulation setup

We generated datasets from two causal topologies where *A* ╨ *C* | *B*: (i) *A* → *B* → *C*, and (ii) *A* ← *B* → *C*. Given a topology, we generated scRNA-seq counts from a Poisson lognormal model dictated by the topology. To test the effect of data sparsity on method performance, we simulated three sparsity levels for gene B. For each of the six conditions (two topologies × three sparsity levels), we generated 100 Monte Carlo replicates, each with 100,000 cells, creating a total of 600 datasets. We detail the precise data-generating process in **Supplementary Note 4**.

#### Statistical inference and performance metrics

For each dataset, we applied ATC-PC and standard partial correlation (Pearson and Spearman on log-transformed counts) to test three conditional independencies: the true one (*A* ╨ *C* | *B*) and the two false ones (*A* ╨ *B* | *C* and *B* ╨ *C* | *A*). We evaluated performance with three metrics: (i) type I error rate, defined as the proportion of replicates where the true conditional independence *A* ╨ *C* | *B* was falsely rejected (*p*-value *<* 0.05); (ii) FDR, defined as the mean false discovery proportion over 100 replicates. For each replicate, discoveries were declared using the Benjamini–Hochberg procedure on all three *p*-values at a significance level of 0.05; and (iii) power, defined as the proportion of hypotheses *A* ╨ *B* | *C* and *B* ╨ *C* | *A* that were correctly rejected over 100 replicates. We report power for both raw and Benjamini–Hochberg adjusted *p*-values at a significance level of 0.05.

### Validating ATC-PC with real data

To validate ATC-PC in a real-data context, we tested its behavior on gold-standard causal motifs constructed from high-confidence experimental evidence. We performed this validation independently on two large-scale perturbation datasets, RPE1 and K562^23,24^, to ensure the robustness of the conclusions.

#### Construction of gold-standard causal motifs

For each dataset, we constructed sets of three-gene causal motifs representing chains (*A* → *B* → *C*), forks (*A* ← *B* → *C*), and colliders (*A* → *B* ← *C*). The edges within these triplets were defined based on high-confidence perturbation effects, filtered stringently by statistical significance, effect size, and the absence of evidence for feedback loops. The resulting motifs were further filtered to ensure their correlation patterns matched theoretical expectations for each topology. The complete, step-by-step procedure is provided in **Supplementary Note 5**.

#### Assessing ATC-PC performance on causal motifs

While the gold-standard motifs were constructed from high-confidence evidence, they existed within a larger, unknown network where other paths can confound the relationships between A, B, and C. For example, for a constructed *A* → *B* → *C* motif, an unknown mediator D can create a parallel path *A* → *D* → *C*. Therefore, rather than performing conditional independence tests, we validated ATC-PC by assessing the “relative decrease” that quantifies the fraction of the original association between two variables *i* and *j* that is explained away by conditioning on *k*:

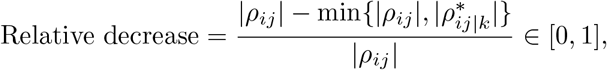

where *ρ*_*ij*_ is the marginal correlation and 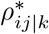 is the partial correlation estimated by ATC-PC. The min 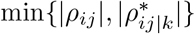 term represents the “causal score” between genes *i* and *j* after conditioning on gene *k*. The three relative decrease values from the gene pairs within a triplet form a statistical signature. We then compared it against the theoretically expected pattern for the motif’s topology:

- For the *A* → *B* → *C* chains, mediator B transmits the causal influence from A to C. Conditioning on B should substantially attenuate the association between A and C, resulting in a large relative decrease. In contrast, we expect smaller decreases for the true edges A–B when conditioning on C and B–C when conditioning on A.
- For the *A* ← *B* → *C* forks, conditioning on a common cause B will also reduce or eliminate the association between A and C, resulting in a large relative decrease. Again, we expect smaller relative decrease for the true edges A–B and B–C.
- For the *A* → *B* ← *C* colliders, A and C are independent causes of a common effect B. Conditioning on B induces association between A and C. Therefore, we expect 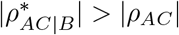, resulting in a relative decrease of at or near zero. For the other pairs, conditioning on an independent cause should also result in a minimal relative decrease.

### Benchmarking GRN inference methods in simulations

#### Synthetic benchmark datasets

We generated ten replicate datasets using the GRN-scPerturbSim framework. We parameterized the simulations using a subset of the large-scale RPE1 dataset, which included all perturbed genes with at least 200 perturbed cells, resulting in a total of 124 genes. This network size is computationally tractable for all benchmarked methods, including computationally intensive algorithms such as GENIE3, GIES, and PC. The ground-truth GRN for each replicate was a scale-free directed network with an average of four regulatory connections per gene. Cell counts for wild-type and perturbation groups were matched to the corresponding RPE1 subsets. The other hyperparameters (see **Supplementary Note 2**) were set as follows: the mean residual variance 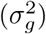 was 1.0, regulatory coefficients (*β*_*gg*_*′* ) were sampled from *U* (*±*[0.3, 0.5]), and the mean perturbation efficacy was 90%.

#### GRN inference methods

The input for all existing methods was log-normalized expression, generated by dividing the counts in each cell by its library size, multiplying by the median library size, and applying a log(*x* + 1) transformation. We compared CausalGRN against seven representative methods (see Extended Data Table 2), grouped by their use of interventional versus observational data.

Methods capable of leveraging interventional data included the score-based causal inference method GIES, which was applied to the full dataset of wild-type and perturbed cells, and Lasso regression applied to the full dataset ignoring cell perturbation labels (Lasso-All).

Methods designed for observational data were applied to wild-type cells only. These included the constraint-based PC algorithm and the score-based GES method. We also benchmarked two tree-based methods, GENIE3 and GRNBoost2, as well as Lasso regression on wild-type cells (Lasso-WT). Implementation details, including specific software packages and key parameters, are elaborated in **Supplementary Note 6**.

#### Performance evaluation

We evaluated the structural accuracy of each inferred GRN against its known ground-truth network. We calculated performance metrics for each of the ten replicate datasets and then averaged the results to ensure robust comparison.

We used four complementary metrics to assess performance. Precision-recall (PR) curves evaluated the overall ranking quality of methods that produced a ranked edge list. We also calculated the F1 score, the Jaccard similarity index, and the Structural Hamming Distance (SHD) on the top *k* predictions (*k* = 1, …, 1000) to specifically quantify the accuracy of the predictions most relevant for experimental validation. The SHD is defined as the number of edge additions, deletions, or reversals required to match the ground-truth graph. For GES and GIES, which return a single unweighted graph rather than a ranked list, we calculated these metrics once on the complete predicted edge set.

### Characterizing and modeling the dominant transcriptional program

Building a complete causal map requires GRN inference methods to account for gene programs. Methods that do not explicitly model these programs risk inferring spurious edges between genes co-regulated by a shared program (Extended Data Fig. 5a). Furthermore, simply regressing out a program’s signature prior to inference is statistically problematic, as conditioning on a common effect can induce spurious correlations among upstream regulators (Extended Data Fig. 5b). Therefore, CausalGRN explicitly models gene programs, beginning with their empirical characterization.

Since gene programs coordinate the expression of downstream targets, their modulation by perturbations should lead to observable coordinated response patterns. Thus, our empirical characterization started by identifying coordinated responses across perturbations. We defined gene responses as binarized (1 if adjusted *p*-value *<* 0.05, 0 otherwise). To focus on robust coordinated signals, we excluded genes unresponsive to any perturbation and, for datasets with few perturbations (e.g., hESC-DE), retained only genes responsive to multiple perturbations. Hierarchical clustering of the filtered genes, based on the Jaccard distance of their binary response profiles, revealed cohesive modules representing dominant coordinated responses (Extended Data Fig. 5c–f).

We next tested if these dominant response modules relate to intrinsic baseline programs present in wild-type cells. We compared the modules to the first principal component calculated from wild-type cells (WT-PC1), which represents the dominant axis of baseline variation and reflects coherent biological processes (Extended Data Fig. 5g). To quantify this link, we summarized each primary module’s response profile using the first principal component of the module’s response significance matrix. We calculated a membership score for all genes by correlating their response significance profile with this module’s response profile. Analysis confirmed a strong association between these membership scores and gene loadings on WT-PC1 (Extended Data Fig. 5h–k), and this strong association validated WT-PC1 as a robust proxy for the dominant response program.

### Incorporating the dominant program into CausalGRN

CausalGRN explicitly models the dominant transcriptional program by incorporating its validated proxy, WT-PC1, into the network structure. First, we calculated the WT-PC1 score for every cell (wild-type and perturbed) and added this score to the expression matrix as an additional “pseudo-gene”, representing the program’s activity.

Next, we defined the set of genes constituting this dominant program. The specific pattern of association observed between WT-PC1 loadings and dominant response module membership (Extended Data Fig. 5h–k) dictated the threshold selection: datasets where strong membership was indicated by both high positive and high negative WT-PC1 loadings (RPE1, K562, and hESC-DE) used a threshold on absolute loading (*>* 0.025); the HCT116 dataset, where strong membership was indicated primarily by high positive loadings, used a threshold on positive loading (*>* 0.02).

CausalGRN was then applied to the data augmented with the WT-PC1 node. Undirected edges were maintained between the WT-PC1 node and the defined program genes to reflect their definitional relationship; the algorithm otherwise proceeded as standard.

### Benchmarking GRN inference methods in real data

To evaluate GRN inference accuracy on real data, we benchmarked all eight methods on the RPE1, K562, HCT116 and hESC-DE datasets. In the absence of a known ground-truth network, we assessed performance using a five-fold cross-validation scheme, where an inferred network’s ability to recapitulate the effects of held-out perturbations serves as a proxy for causal accuracy.

#### Cross-validation and network inference strategy

To construct a benchmark that balances statistical power with computational feasibility, we first curated a subset of high-quality perturbations from each dataset. We filtered each dataset for perturbations with a sufficient number of cells to reliably measure perturbation effects, while limiting the total number of perturbations for computational tractability. This resulted in 267 knockdowns with at least 150 perturbed cells for the RPE1 dataset, 159 knockdowns with at least 400 perturbed cells for the K562 dataset, 199 knockdowns with at least 300 perturbed cells for the HCT116 dataset (with wild-type cells downsampled to 50,000), and all 14 knockouts from the hESC-DE dataset. These curated sets of perturbations were then randomly partitioned into five folds to serve as the held-out test sets.

We defined the network gene set for each dataset to ensure computational tractability, which is primarily dictated by network size. For the large-scale RPE1, K562 and HCT116 datasets, the network was restricted to the curated knockdown genes. For the smaller-scale hESC-DE dataset, the network gene set was constructed from the 14 knockout genes and a randomly selected subset of their strongly correlated genes (absolute Pearson correlation *>* 0.05), resulting in a final, unique set of 152 genes.

We then applied all eight algorithms, tailoring each to its design. Methods capable of leveraging interventional data were trained on the combined dataset of wild-type cells and perturbed cells from the training folds. In contrast, methods designed for observational data were trained only on the wild-type cells. Specific implementation details and parameters for each method are elaborated in **Supplementary Note 6**.

#### Evaluation of inferred edges using a perturbation-effect proxy

Evaluating the accuracy of an inferred GRN on real data is challenging due to the absence of a comprehensive ground-truth network. We therefore devised an evaluation strategy based on the principle that a high-confidence inferred causal edge (e.g., *A* → *B*) should correspond to a significant, experimentally observable change in the expression of gene B when gene A is perturbed. We used the magnitude of this downstream effect (quantified by absolute Cohen’s D) as a robust proxy for edge validity.

We implemented this evaluation within the five-fold cross-validation. In each fold, we inferred a GRN using the training data and identified inferred edges whose source genes were held-out perturbation targets. For each of these candidate edges, we then calculated its causal effect proxy: the absolute Cohen’s D of the target gene’s expression change in the held-out perturbation experiment. CausalGRN infers a network including the WT-PC1 pseudo-gene, a feature not present in the other methods. To enable a direct and fair comparison, we projected the CausalGRN output into the same gene-only space as the other methods. This was achieved by converting all edges of the form *A* → WT-PC1 into directed edges from A to each of the program genes, and then removing the WT-PC1 pseudo-gene node.

To generate the final performance curves, we pooled all candidate edges evaluated on the test sets from the five folds and ranked them by the confidence scores assigned by the inference method (see Extended Data Fig. 3 for an example). The final metric, which we term “CV mean |Cohen’s D|”, is the mean of the proxy effect sizes for the top *N* edges in this global ranking, plotted as a function of *N* . This metric provides a direct measure of whether an algorithm’s high-confidence predictions correspond to strong causal effects. For methods that return an unweighted set of edges (GES and GIES) rather than a ranked list, performance was assessed by a single calculation of this metric on the complete predicted edge set.

### Benchmarking perturbation effect prediction in simulations

#### Simulation dataset and cross-validation

We benchmarked our framework using replicate 1 of the GRN-scPerturbSim synthetic dataset, which was also used for the GRN inference evaluation. To assess the models’ ability to generalize to unseen perturbations, we implemented a five-fold cross-validation by randomly partitioning the perturbation experiments.

#### Benchmark frameworks

We evaluated our framework against two baselines using this identical cross-validation scheme. The three frameworks were as follows.

- CausalGRN-guided: For each cross-validation split, a GRN was first inferred from the training data. This framework builds expression models using linear regression on the inferred causal parents and applies our perturbation effect propagation algorithm.
- Average Effect: a simple baseline that predicts the training-set average perturbation effect for all test perturbations.
- Complete Graph-guided: a non-causal baseline. This framework builds expression models using Lasso regression on all other genes and applies the same propagation algorithm.

#### Oracle analysis

We performed an ablation study using an “oracle” setting to isolate the performance of the fitted expression models from the challenge of effect propagation. In this analysis, instead of propagating effects from the initial perturbation, the fitted models (both CausalGRN-guided and Complete Graph-guided) were applied directly to the true, observed average expression levels of their respective predictor genes from the held-out test cells. This analysis pinpoints whether a framework’s failure is due to poor underlying expression models or a failure during the propagation step.

#### Evaluation of DEG prediction accuracy

We evaluated performance using the precision of predicted DEGs from held-out perturbations. Predictions from all five cross-validation folds were pooled to generate aggregate performance curves, after excluding all self-effects (the effect of perturbing a gene on itself). We generated separate precision curves for down- and up-regulated DEG prediction. Precision was calculated as the fraction of true down- (or up-) regulated DEGs found within the top *N* genes ranked by the most negative (or positive) predicted differential expression, respectively.

### Benchmarking perturbation effect prediction in real data

#### Datasets and cross-validation

We benchmarked perturbation effect prediction on the three large-scale datasets (RPE1, K562 and HCT116) using a five-fold crossvalidation. This analysis used the identical curated perturbation subsets from the GRN inference benchmark.

#### Benchmark frameworks

We evaluated our framework against the Average Effect and Complete Graph-guided baselines. We implemented these baselines identically to the simulation benchmark for perturbation effect prediction, operating on the original gene set without the WT-PC1 program node.

The CausalGRN-guided framework explicitly models the WT-PC1 program. This required a specialized prediction strategy for that node, built to leverage our observation that a perturbation’s effect on this program can be portable across different datasets. We therefore built a portability model specifically for the WT-PC1 node. This linear model was trained to predict the WT-PC1 response in a target dataset. It used as predictors the observed response of the target dataset’s WT-PC1 program to the same perturbation, as measured in the reference datasets (e.g., using K562 and HCT116 as references when RPE1 was the target).

To predict the effect of a test perturbation, the framework first established the initial signal. This signal combined two components: (1) the direct effect on the perturbed gene (i.e., its perturbed expression level) and (2) the response of the WT-PC1 program. To determine the program’s response, the framework applied the trained portability model. If a program-specific response could not be confidently predicted (i.e., the portability model was unavailable for the test perturbation or failed the Fstatistic significance threshold of *p*-value *<* 0.1), the framework instead substituted the training-set average effect for both the WT-PC1 node and its program genes. This complete initial signal was then propagated through the causal network to predict the final, system-wide gene expression changes.

#### Oracle analysis and evaluation

We also evaluated the CausalGRN-guided and Complete Graph-guided frameworks in the “oracle” setting, as described previously. All frameworks were evaluated using the identical DEG prediction accuracy metrics from the simulation benchmark for perturbation effect prediction.

### Application to the hESC-beta dataset

#### Dissecting the upstream regulation of Type 2 Diabetes (T2D) risk genes

We applied CausalGRN to 327 genes, defined as the union of 312 T2D risk genes from Liu et al. ^27^ with expression sparsity *<* 90% in the hESC-beta dataset, the 8 knockout targets, and 8 additional developmental genes. We included WT-PC1 (derived from all genes in the dataset) as a “pseudo-gene” in the network to represent the primary gene program and defined its program genes as those with absolute WT-PC1 loadings *>* 0.025. For comparison, we also constructed a correlation network on the same gene set by applying the WGCNA approach ^28^ to all cells.

#### Prioritizing upstream regulators of T2D risk genes

We prioritized upstream regulators of T2D risk genes using a descendant enrichment analysis. For each T2D risk gene, we selected one control gene with the closest average expression among all other genes in the dataset, yielding an expression-matched control gene set. We then added these control genes to the CausalGRN analysis so that each knockout gene’s downstream neighborhood could be tested for enrichment of T2D risk genes relative to expression-matched controls. For each knockout gene, we defined its downstream set as all descendants within a maximum directed path length of two, treating undirected edges as bidirectionally traversable. We then quantified enrichment of T2D risk genes among these descendants relative to the matched controls using an odds ratio and assessed significance using a two-sided Fisher’s exact test. *P* -values were adjusted using the Benjamini-Hochberg procedure.

## Supporting information

Supplementary Algorithm 1 and Supplementary Notes 1-6

## Supplementary information

Supplementary Algorithm 1 and Supplementary Notes 1–6.

### Acknowledgements

This work was supported by NIH grant U01HG013177.

## Data availability

All datasets analyzed in this study are publicly available. The RPE1 and K562 Perturb-seq datasets ^24^ were downloaded from Figshare (https://doi.org/10.25452/figshare.plus.20029387.v1). The HCT116 dataset ^26^ was downloaded from Figshare (https://doi.org/10.25452/figshare.plus.29190726.v1). The hESC-DE and hESC-beta datasets ^27^ were downloaded from the Gene Expression Omnibus (GEO) under accession number GSE313516 (https://www.ncbi.nlm.nih.gov/geo/query/acc.cgi?acc=GSE313516).

## Code availability

The CausalGRN package is available at https://github.com/yub-hutch/CausalGRN.

## Author contributions

B.Y. and W.S. conceived the study and designed the computational framework. B.Y. implemented the software, performed all analyses, and drafted the manuscript. G.Q., L.H., and A.S. provided regular guidance on the study and revised the manuscript; A.S. contributed to methodological development. D.L. and D.H. provided expertise on the hESC-DE and hESC-beta datasets and contributed to the biological interpretation of the results. W.S. supervised the study and substantially revised the manuscript. All authors approved the final manuscript.

## Extended Data

**Extended Data Fig. 1.**
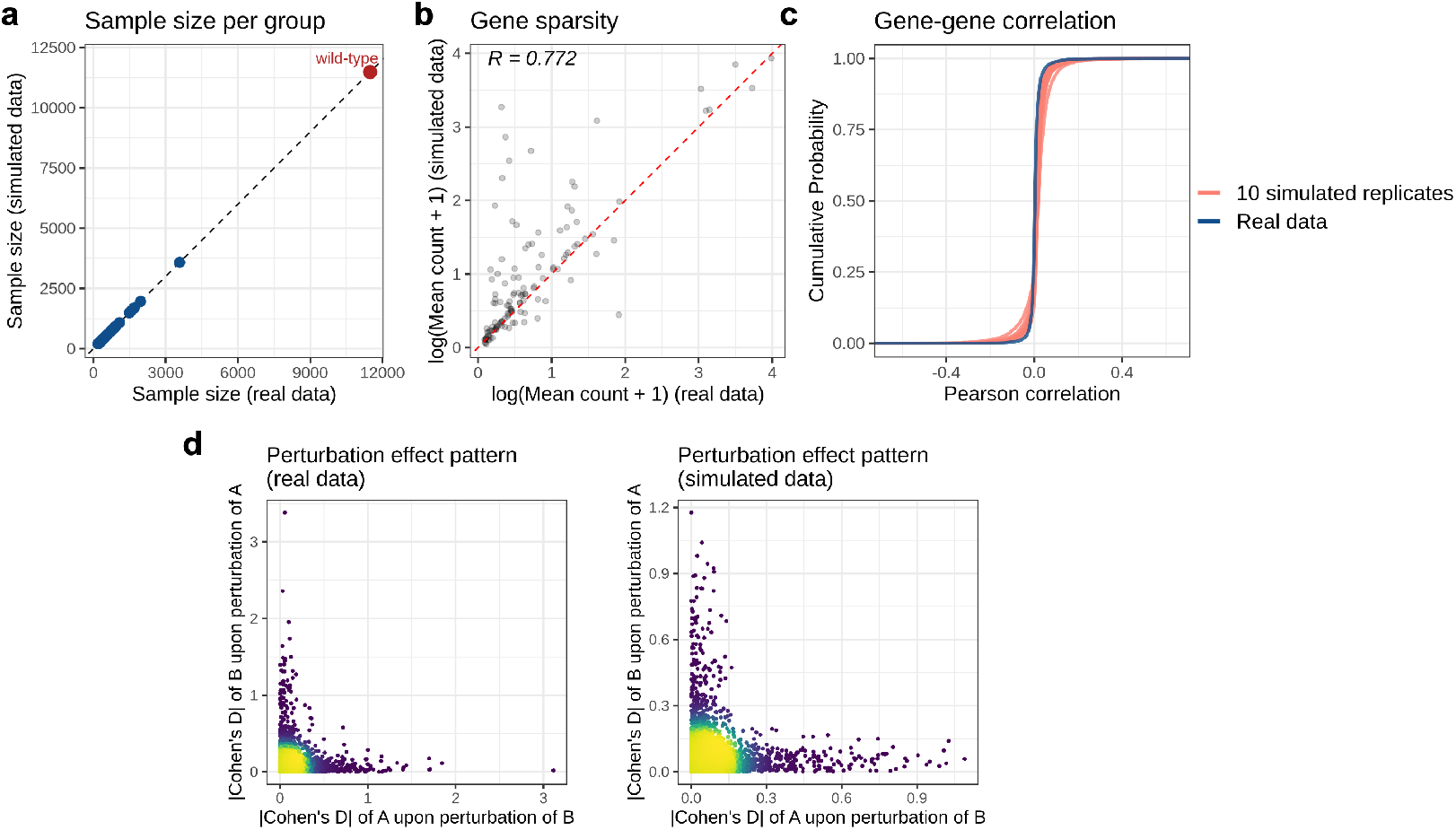
GRN-scPerturbSim recapitulates key statistical properties of the target RPE1 dataset subset. Comparisons between a target RPE1 dataset subset and ten simulated replicates by GRN-scPerturbSim. The target dataset includes 124 genes that were individually perturbed (see Methods). **a**, Sample size per group. Cell counts for wild-type (red point) and each of the 124 perturbation groups were matched to the target dataset across ten replicates by design. **b**, Gene sparsity in wild-type cells. Each point represents the log-transformed mean count of a gene in the target dataset versus replicate 1. Pearson’s *R* = 0.772 for replicate 1; mean *R* = 0.804, s.d. = 0.045 across ten replicates. **c**, Gene-gene correlation distribution. Empirical cumulative distribution functions of gene-gene correlations in target dataset (covariates adjusted; see Methods) versus ten replicates (log-transformed). **d**, Perturbation effect pattern. For each gene pair A–B, the plots show the effect (absolute Cohen’s D) of perturbing B on A’s expression against the effect of perturbing A on B’s expression, in the target dataset versus replicate 1.

**Extended Data Fig. 2.**
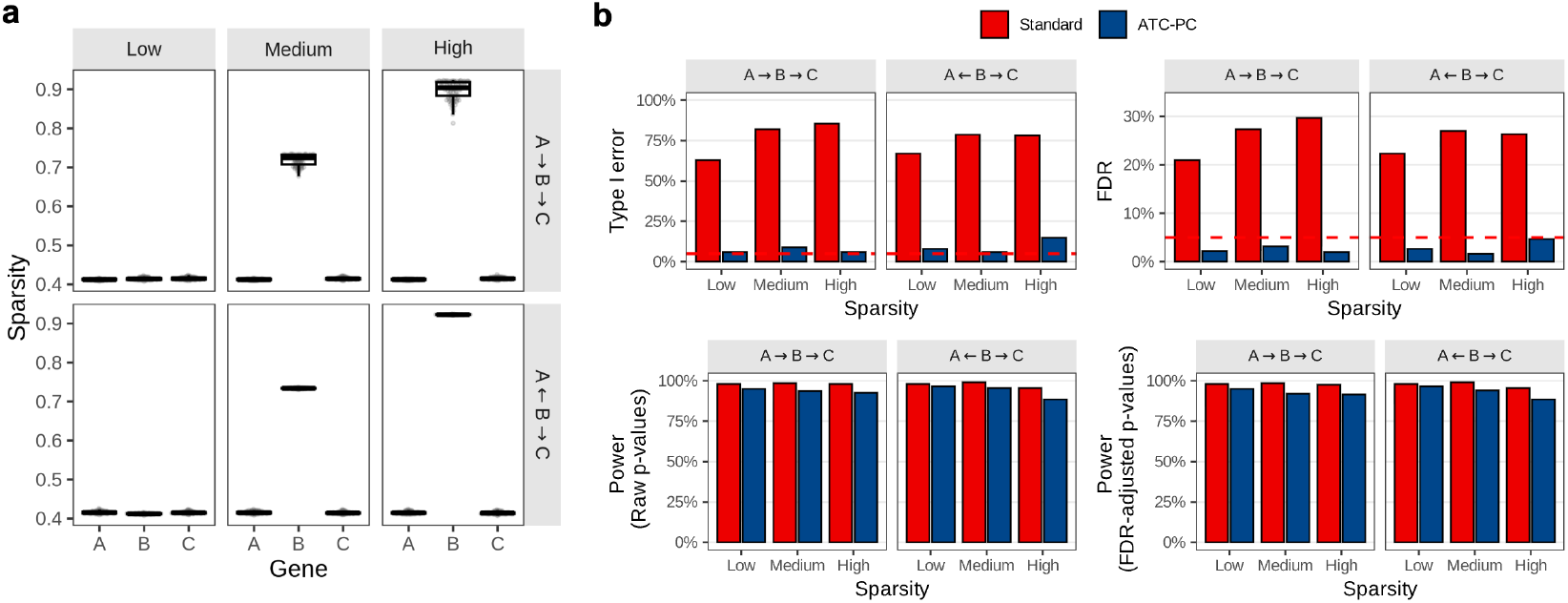
Simulation details and validation of ATC-PC using Spearman correlation. **a**, Sparsity levels for each gene in the two simulated causal topologies (*A* → *B* → *C* and *A* ← *B* → *C*). **b**, Performance of ATC-PC compared to standard partial correlation, replicating the analysis from Fig. 2a–d but using the non-parametric Spearman correlation. Panels show Type I error and FDR for the spurious A–C edge, and statistical power for the true A–B and B–C edges. The dashed red line indicates the nominal 0.05 significance level. All metrics were calculated from 100 simulation replicates.

**Extended Data Fig. 3.**
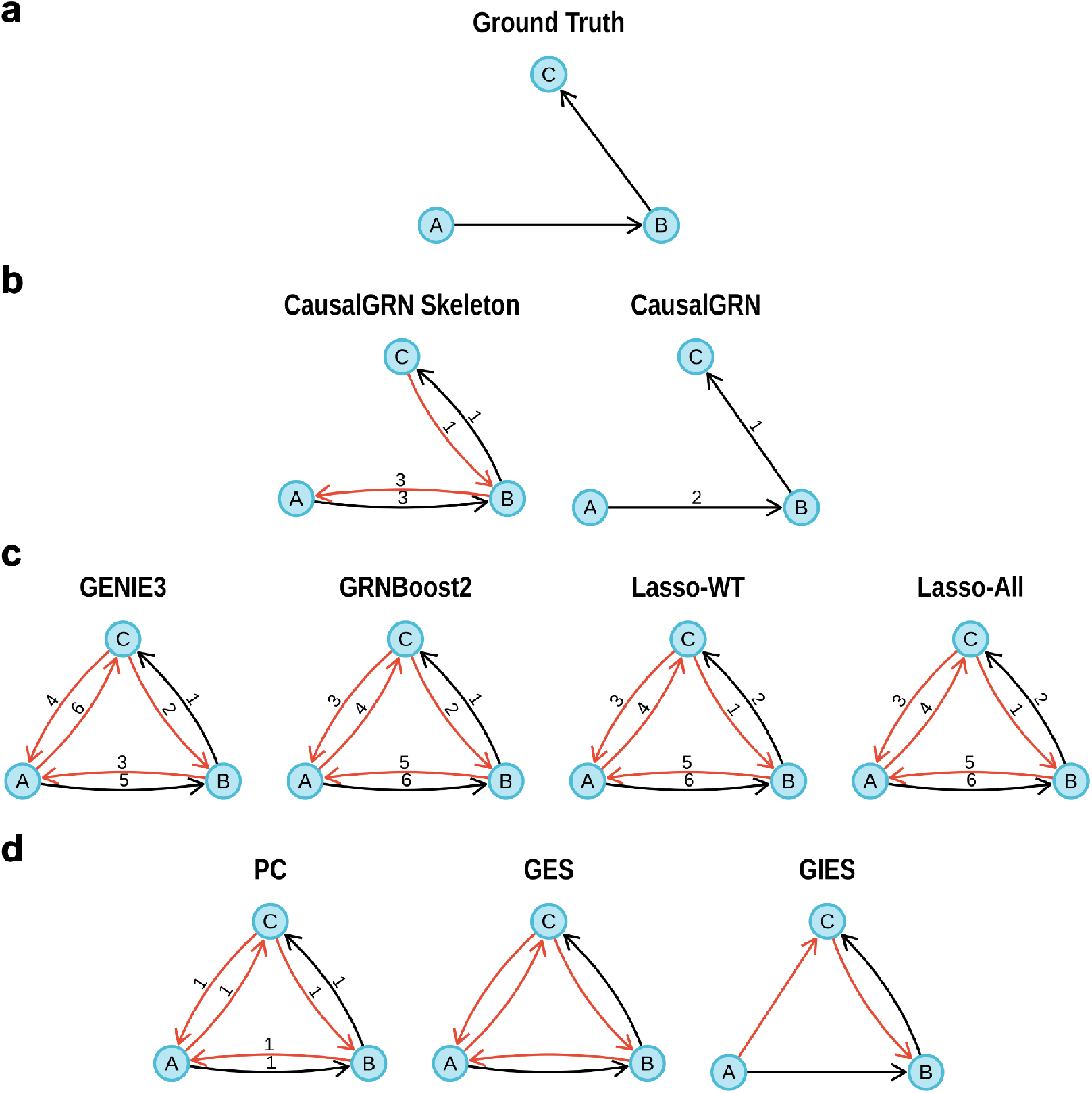
CausalGRN identifies a simple causal chain that existing methods fail to. Data were simulated for 10^5^ wild-type and 10^4^ A-perturbed cells from a chain *A* → *B* → *C* (**a**). Latent log-expressions were generated as *u*_*A*_ ∼ 𝒩 (0, *σ*^2^), *u*_*B*_ ∼ 𝒩 (*u*_*A*_ + *s, σ*^2^), and *u*_*C*_ ∼ 𝒩 (*u*_*B*_ − *s, σ*^2^) with *σ* = 2 and *s* = −4. Counts were drawn from Poisson(exp(*u*_*g*_)) for *g* ∈ {*A, B, C*}. The *s* = −4 intercept induces 86.4% zero counts for gene B. Perturbed cells were generated by introducing a 90% knockdown of gene A, applied as an additive shift of log(1 − 0.9) to *u*_*A*_. This structure creates a classic challenge: A and C are conditionally independent given B yet exhibit a strong marginal correlation (Pearson’s *R* = 0.49 on log-transformed counts). The inferred GRNs for methods grouped by category (see Extended Data Table 2) are shown in panels **b–d**. Edges are colored **black** if they match ground truth and **red** if they are incorrect. **Edge labels indicate each method’s rank for the edge (1=highest confidence)**; GES and GIES output graphs without edge ranks. **a**, The ground-truth network *A* → *B* → *C*. **b**, CausalGRN first identifies the correct skeleton (left) by recovering the conditional independence between A and C given B, then uses the perturbation of A to orient all edges (right). **c**, Outputs from association-based methods. Lasso-WT: Lasso applied to wild-type cells; Lasso-All: Lasso applied to all cells (wild-type and perturbed). All methods fail to distinguish causality from mere correlation. **d**, Outputs from classical causal inference methods. They fail to find the correct topology as their models are unsuitable for sparse data.

**Extended Data Fig. 4.**
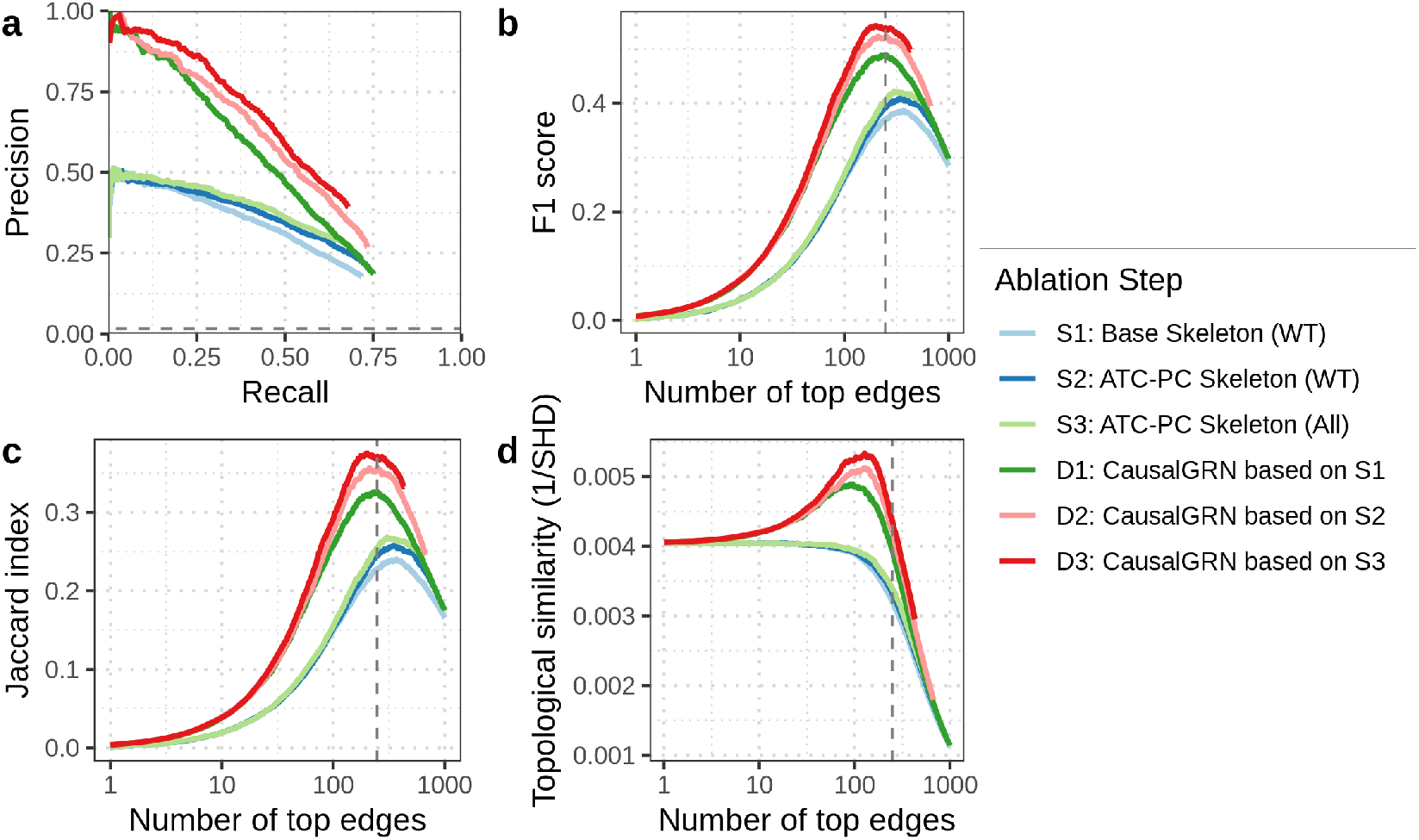
CausalGRN’s performance is driven by synergistic contributions from its two-stage design. This ablation study dissects the contributions of CausalGRN’s components on the simulated benchmark (averaged over ten replicates). We evaluated the performance of the undirected skeleton (S-curves) and the final directed graph (D-curves) across four metrics: **a**, Precision-Recall; **b**, F1 score; **c**, Jaccard index; and **d**, Topological similarity (1/SHD). The horizontal dashed line in **a** indicates random precision. The vertical dashed lines in **b–d** indicate the average number of true edges in the ground-truth networks. The configurations are: S1 (Skeleton with standard partial correlation, on wild-type cells), S2 (Skeleton with ATC-PC, on wild-type cells), S3 (Skeleton with ATC-PC, on all cells), D1 (CausalGRN orientation based on S1), D2 (CausalGRN orientation based on S2), and D3 (CausalGRN orientation based on S3). The results demonstrate that all three components of the CausalGRN design provide cumulative contributions to the final performance: the ATC-PC (e.g., S1 vs S2), the use of all cells for skeleton inference (e.g., S2 vs S3), and the perturbation-guided orientation (e.g., S3 vs D3).

**Extended Data Fig. 5.**
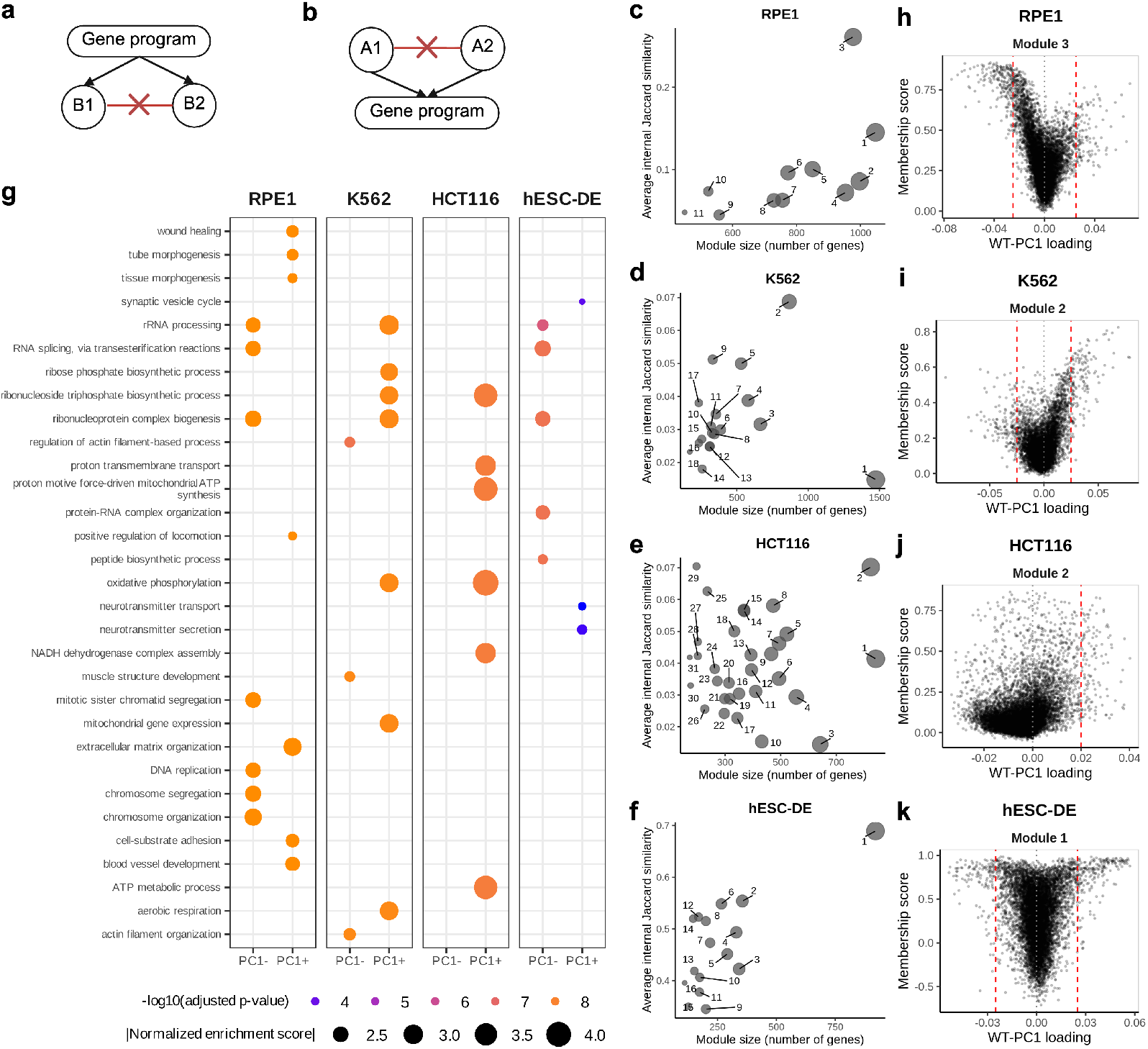
WT-PC1 serves as a proxy for the dominant response program. **a, b**, Schematics illustrating potential inference errors. Black edges show true relationships; red edges marked with ‘X’ show spurious links incorrectly inferred (**a**) between genes co-regulated by a shared program when ignoring the program, or (**b**) between upstream regulators when regressing out their common effect (the program). **c–f**, Coordinated response modules across perturbations in RPE1 (**c**), K562 (**d**), HCT116 (**e**), and hESC-DE (**f** ) datasets. We identified these modules by clustering gene response profiles using Jaccard similarity on binarized responses. Plots show module size versus average internal Jaccard similarity; point size is proportional to module size, highlighting dominant response modules as those with both large size and high cohesion. **g**, WT-PC1 is associated with specific biological processes across datasets. Gene Set Enrichment Analysis dot plot shows significantly enriched (adjusted *p*-value *<* 0.01), simplified (similarity cutoff = 0.5) Gene Ontology Biological Process terms associated with positive (PC1+) and negative (PC1-) WT-PC1 loadings. For each sign and dataset, the top 20 most significant terms were selected, and then filtered to show only those with an absolute normalized enrichment score ≥ 2 (no terms met criteria for HCT116 PC1-). Color reflects the enrichment adjusted *p*-value; dot size reflects the magnitude of normalized enrichment score. **h–k**, WT-PC1 loadings reflect participation in dominant response modules across datasets. Scatter plots show gene loadings on WT-PC1 versus membership score (correlation between a gene’s response significance profile and the primary module’s representative response profile) for the primary module identified in c–f. These plots confirm the strong association and visualize the specific patterns that empirically guided the selection of the WT-PC1 loading thresholds (red dashed lines) used by CausalGRN (see Methods).

**Extended Data Fig. 6.**
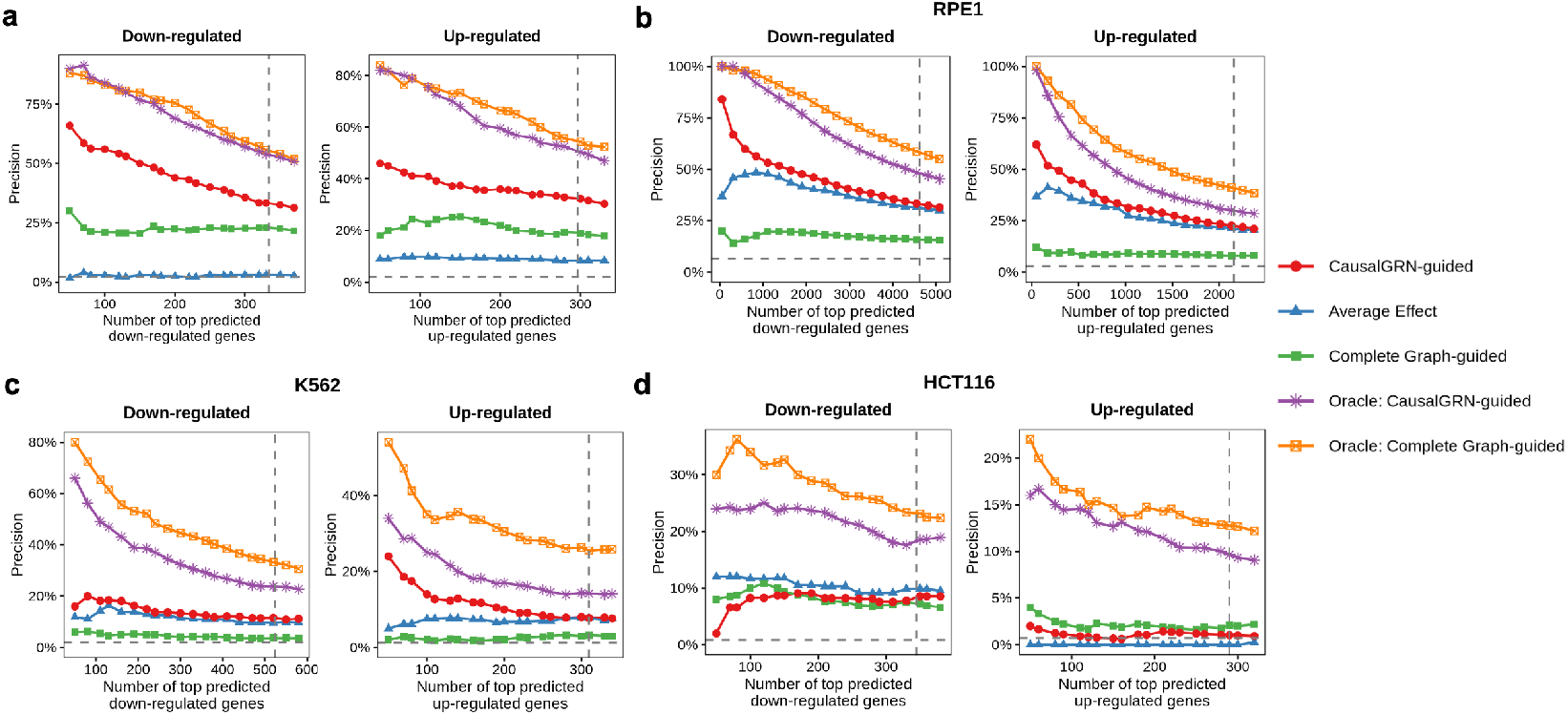
Oracle analysis of perturbation effect prediction frameworks. Precision in identifying true down-regulated (left column) and up-regulated (right column) DEGs from held-out perturbations in five-fold cross-validation. Predictions from all folds were pooled to generate each curve. Panels show results for (**a**) the GRN-scPerturbSim simulated dataset, (**b**) RPE1, (**c**) K562, and (**d**) HCT116 datasets. Precision (y-axis) is the fraction of true down- (or up-) regulated DEGs found within the top *N* genes (x-axis) ranked by the most negative (or positive) predicted differential expression, respectively. Performance is compared for the Average Effect baseline, which predicts the training-set average perturbation effect for all test perturbations, and two model-building strategies. The CausalGRN-guided strategy (models built with linear regression on inferred causal parents) is shown in its practical, propagation-based form (see Methods) and its oracle version. The Complete Graph-guided strategy (models built with Lasso regression on all other genes) is similarly shown in its propagation-based and oracle versions. The oracle versions establish the theoretical upper-bound performance for each strategy by applying the fitted models to the true, observed average expression levels of their respective predictors from the test cells. The horizontal dashed line indicates random precision; the vertical dashed line marks the total number of true down- or up-regulated DEGs, respectively, in the pooled held-out test set.

**Extended Data Fig. 7.**
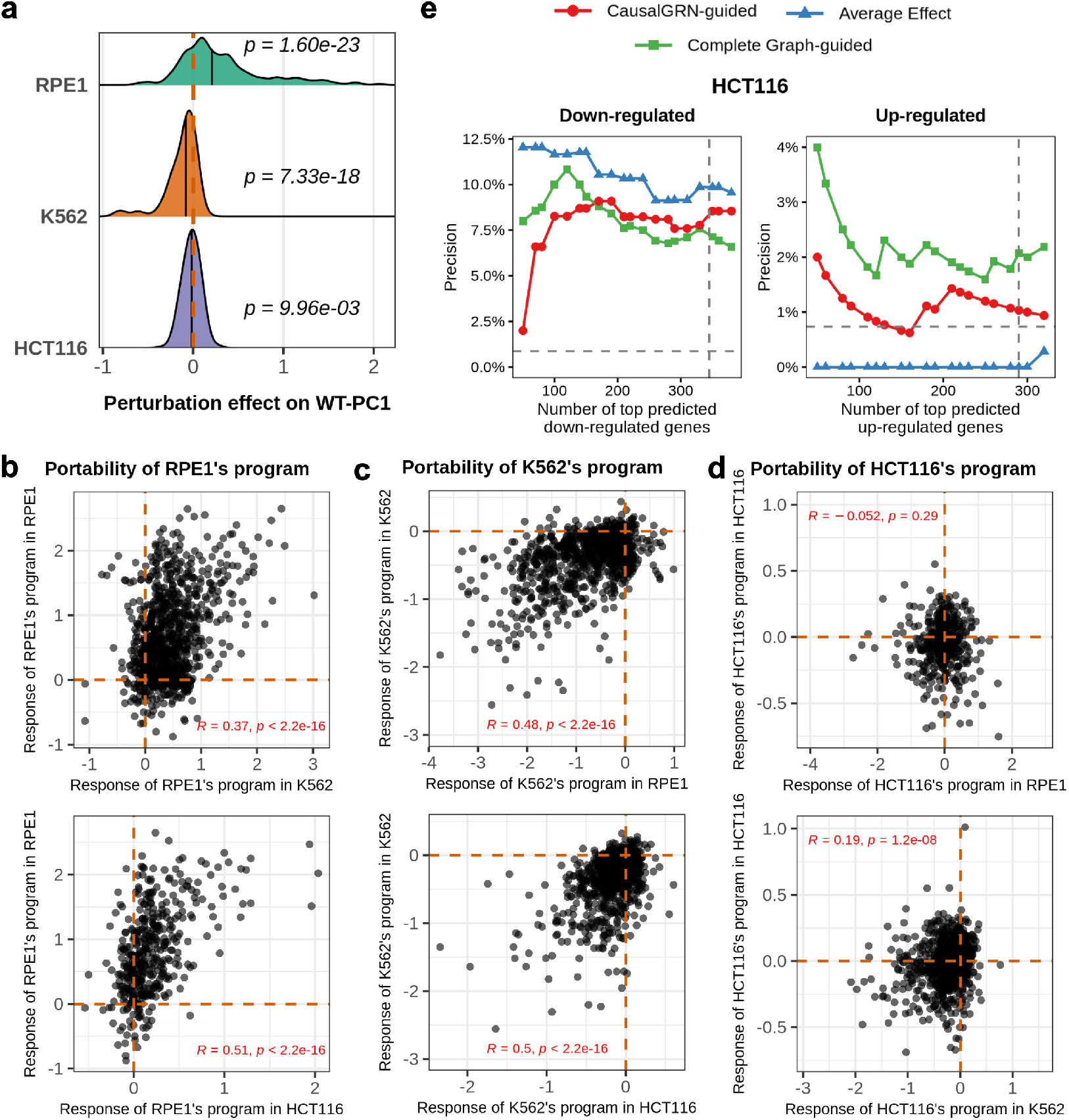
Perturbation effect prediction accuracy in HCT116 and portability of gene program responses across cell lines. **a**, Ridged density plot showing that many perturbations alter the WT-PC1 program in the same direction. Each distribution represents the perturbation effects on WT-PC1 in a given dataset (RPE1, K562, HCT116). A perturbation’s effect (x-axis) is the difference in mean WT-PC1 program score between its perturbed cells and wild-type cells. *P* -values (one-sample Wilcoxon signed-rank test versus 0) confirm the median of each distribution is significantly non-zero. **b–d**, Portability of WT-PC1 program responses across datasets. Each scatter plot compares a program’s response to a perturbation (Cohen’s d) in its source dataset (y-axis) against its response to the same perturbation in a reference dataset (x-axis). Pearson correlation coefficients with *p*-values are shown. **e**, Precision in identifying true down-regulated (left column) and up-regulated (right column) DEGs from held-out perturbations in five-fold cross-validation in the HCT116 dataset. Precision (y-axis) is the fraction of true down- (or up-) regulated DEGs found within the top *N* genes (x-axis) ranked by the most negative (or positive) predicted differential expression, respectively. The plot compares three prediction frameworks: CausalGRN-guided, Average Effect, and Complete Graph-guided. The horizontal dashed line indicates random precision; the vertical dashed line marks the total number of true down- or up-regulated DEGs, respectively, in the pooled held-out test set.

**Extended Data Table 1.** Comparison of scRNA-seq data simulators. Splatter ^38^ and SPARSim ^39^ reproduce dataset statistics without imposing a known underlying GRN. SERGIO ^40^ and dyngen ^41^ simulate GRN-driven dynamics but are not designed to be directly parameterized to match a target dataset. GeneSPIDER2 ^42^ uses generically parameterized models rather than learning from a target dataset. GRouNdGAN ^43^ assumes a bipartite transcription factor (TF) → target gene structure, limiting TF-TF interactions and multi-layer cascades.

| Simulator | Supports<br>complex GRN | Simulates<br>perturbations | Mimics<br>target dataset | Models<br>heterogeneous<br>perturbation efficacy |
| --- | --- | --- | --- | --- |
| GRN-scPerturbSim | ✓ | ✓ | ✓ | ✓ |
| *Data-driven simulators* |  |  |  |  |
| Splatter | ✗ | ✗ | ✓ | ✗ |
| SPARSim | ✗ | ✗ | ✓ | ✗ |
| *GRN-driven trajectory simulators* |  |  |  |  |
| SERGIO | ✓ | ✓ | ✗ | ✓ |
| dyngen | ✓ | ✓ | ✗ | ✓ |
| *GRN-driven steady-state simulators* |  |  |  |  |
| GeneSPIDER2 | ✓ | ✓ | ✗ | ✓ |
| GRouNdGAN | ✗ | ✓ | ✓ | ✗ |

**Extended Data Table 2.**
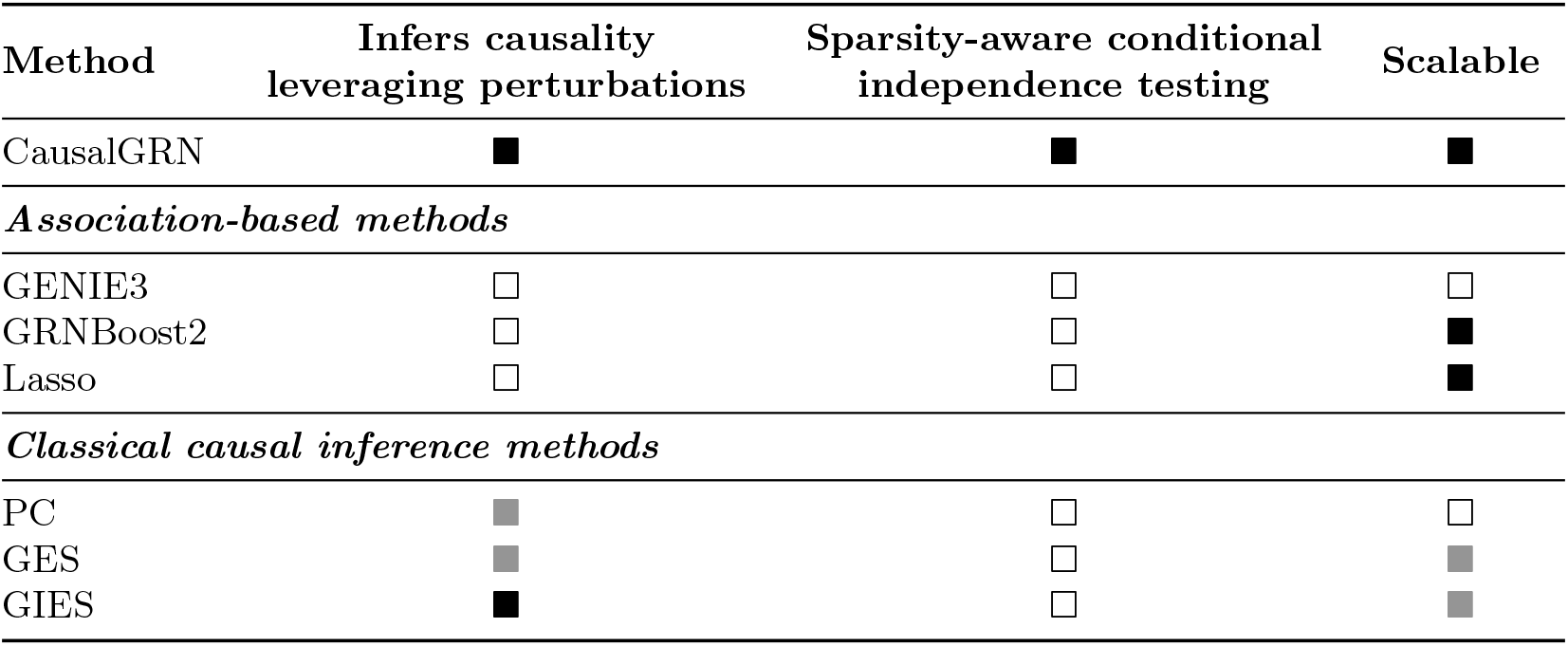
Comparison of CausalGRN with existing GRN inference methods. Ratings of methods are indicated by squares (■ = strong; ■ = moderate; □ = not available). Association-based methods GENIE3 ^3^, GRNBoost2 ^4^ and Lasso do not infer causality. Classical causal inference methods PC ^6^, GES ^7^ and GIES ^8^ are unsuitable for sparse scRNA-seq data and cannot scale computationally to genome-scale networks.

