## Supplementary Algorithm 1 and Supplementary Notes 1-6 for "CausalGRN: deciphering causal gene regulatory networks from single-cell CRISPR screens"

### Supplementary Information for "CausalGRN: deciphering causal gene regulatory networks from single-cell CRISPR screens"

#### Contents

**Supplementary Algorithm 1: Causal inference of GRN skeleton** **2**

**Supplementary Note 1: Transitive orientation of CausalGRN** **3**

**Supplementary Note 2: GRN-guided simulation of single-cell perturbation data (GRN-scPerturbSim)** **4**

**Supplementary Note 3: Perturbation effect prediction via network propagation** **6**

**Supplementary Note 4: Simulation details for ATC-PC validation** **8**

**Supplementary Note 5: Construction of gold-standard causal motifs** **9**

**Supplementary Note 6: Implementation details of GRN inference methods** **10**

### Supplementary Algorithm 1: Causal inference of GRN skeleton

**Algorithm 1** Causal inference of GRN skeleton

**Require:** Complete graph $\mathcal{G}^0 = (V, E^0)$, maximum order of conditional independence test $L = 1$, significance level $\alpha$, threshold of absolute partial correlation $\rho_0$

**Ensure:** Inferred skeleton $\mathcal{G}^{\text{skeleton}} = (V, E^{\text{skeleton}})$, separation set $S(i, j)$ for $i - j \notin E^{\text{skeleton}}$, edge scores $|\rho_{ij}|^{\min}$

$\mathcal{G}^{\text{skeleton}} \Leftarrow \mathcal{G}^0$
$|\rho_{ij}|^{\min} \Leftarrow \infty$ for all edges $i - j$ in $E^0$
$l \Leftarrow 0$
**while** $l \leq L$ and $E^{\text{skeleton}}$ is not empty **do**
    $\mathcal{G}^{\text{working}} \Leftarrow \mathcal{G}^{\text{skeleton}}$
    **for** edge $i - j \in E^{\text{working}}$ (in parallel) **do**
        $N_i \Leftarrow$ neighbors of $i$ in $\mathcal{G}^{\text{working}}$
        $N_j \Leftarrow$ neighbors of $j$ in $\mathcal{G}^{\text{working}}$
        $\mathcal{A}_{ij} \Leftarrow (N_i \cup N_j) \setminus \{i, j\}$
        **for** $\mathcal{C} \subseteq \mathcal{A}_{ij}$ and $|\mathcal{C}| = l$ **do**
            Test $i \perp\!\!\!\perp j \mid \mathcal{C}$ using the ATC-PC method to get partial correlation estimate $\rho_{ij|\mathcal{C}}$ and $p$-value $p_{ij|\mathcal{C}}$
            $|\rho_{ij}|^{\min} \Leftarrow \min(|\rho_{ij}|^{\min}, |\rho_{ij|\mathcal{C}}|)$
            **if** $p_{ij|\mathcal{C}} > \alpha$ or $|\rho_{ij|\mathcal{C}}| < \rho_0$ **then**
                $E^{\text{skeleton}} \Leftarrow E^{\text{skeleton}} \setminus \{\text{edge } i - j\}$
                $S(i, j) \Leftarrow \mathcal{C}$
                **break**
            **end if**
        **end for**
    **end for**
    $l \Leftarrow l + 1$
**end while**

### Supplementary Note 1: Transitive orientation of CausalGRN

This note details the optional transitive orientation procedure, which can be applied after direct orientation to increase the number of directed edges in the final network. It represents a trade-off between the density of oriented edges and the higher confidence of direct orientation, and is particularly useful when the number of perturbed genes is limited.

The procedure uses the set of directly oriented edges to resolve more distant, undirected edges. For a path $A \to B - C$, where A is the perturbed gene, the edge B–C is oriented as $B \to C$ if:

- B was not itself perturbed,
- C is a DEG upon perturbation of A, and
- no other directed path from A to C bypasses B, where an undirected edge is considered traversable in both directions.

To ensure a deterministic and conservative final network, all such candidate orientations are identified across all perturbation experiments before being applied simultaneously. Any edge with conflicting directional evidence arising from this process is left undirected.

### Supplementary Note 2: GRN-guided simulation of single-cell perturbation data (GRN-scPerturbSim)

Given a target single-cell perturbation dataset, GRN-scPerturbSim generates a synthetic dataset mimicking its statistical properties with a known underlying GRN in four steps.

##### *Step 1: Ground-truth network generation*

The simulation is guided by a ground-truth GRN generated using the Barabási-Albert model to produce a scale-free topology characteristic of biological networks [1, 2].

##### *Step 2: Model Parameterization*

Gene expression is modeled using a Poisson lognormal distribution dictated by the generated GRN. The observed count $Y_{ig}$ for gene $g$ in cell $i$ is sampled from $\text{Poisson}(s_i \exp(u_{ig}))$, where $s_i$ is the cell-specific size factor and $u_{ig}$ is the latent log-expression. This latent variable is defined by a linear structural equation based on the GRN:

$$u_{ig} \sim \mathcal{N}(\beta_{g0} + \sum_{g' \in \text{Pa}_g} \beta_{gg'} u_{ig'}, \sigma_g^2),$$

where $\text{Pa}_g$ are the parent regulators of gene $g$ in the GRN. The model parameters are derived from the target perturbation dataset to ensure quantitative realism. Specifically, the cell-specific size factors ($s_i$) preserve empirical cell-to-cell variation in sequencing depth from the target dataset. The basal log-expression levels ($\beta_{g0}$) are set to match the empirical mean levels from the target dataset. The residual gene-specific variances ($\sigma_g^2$) are drawn from a Gamma distribution to establish a baseline noise level. Finally, the regulatory coefficients ($\beta_{gg'}$) are sampled from a uniform distribution with randomized signs; the bounds of this distribution are calibrated to ensure the resulting gene-gene correlation distribution approximates that of the target dataset.

##### *Step 3: Generating wild-type cells*

Expression of wild-type cells is sampled from the parameterized model. The sampling follows a topological gene order of the GRN, ensuring that the expressions of all parent genes are generated before being used to simulate the expression of a child gene.

##### *Step 4: Generating perturbed cells*

The cellular response to a genetic perturbation is simulated as a two-stage process: a direct effect on the target gene, followed by indirect effects that cascade through the network.

The direct effect is modeled to reflect that CRISPR-based perturbations are rarely fully penetrant and can vary from cell to cell. The simulation assigns each perturbed cell $i$ a “perturbation efficacy”, $e_i$, drawn from a Beta distribution. This $e_i$ represents the fractional reduction in target gene expression. The latent log-expression of the target gene, $u_{ig}$, is then reduced to a new value corresponding to this efficacy, allowing the perturbation to propagate through the network.

The indirect effects are an emergent property of the simulation, as the initial change to the target gene naturally propagates through the network to its descendants. This process generates a complex and biologically plausible pattern of downstream expression changes.

### Supplementary Note 3: Perturbation effect prediction via network propagation

This note provides the complete mathematical description of the CausalGRN perturbation effect prediction framework, which propagates the effect of an initial perturbation through a linear model of GRN.

We assume that at steady state, gene expression is governed by the linear system:

$$X = \alpha + BX + e,$$

where $X$ is the $G \times 1$ vector of gene expression levels for $G$ genes, $\alpha$ is a vector of intercept terms, $B$ is the $G \times G$ regulatory coefficient matrix, and $e$ is a noise vector.

#### Training of expression prediction models

We estimate the parameters $\alpha$ and $B$ from the available single-cell data. The matrix $B$ is sparse, with its structure constrained by the inferred causal GRN.

For each gene $i$, we fit a linear regression model to estimate its endogenous regulation. The predictors for $x_i$ are the expression levels of its parent set, $\mathrm{Pa}(i)$, as defined by the inferred causal GRN. This parent set includes all genes $j$ such that there is a directed edge ($j \to i$) or a bidirectionally-traversable undirected edge ($j - i$) connecting to $i$ in the network.

The model for gene $i$ is:

$$x_i = \alpha_i + \sum_{j \in \mathrm{Pa}(i)} B_{ij} x_j + e_i.$$

To learn the endogenous regulatory function, this model is trained using all cells except those in which gene $i$ itself was directly perturbed.

Collecting the fitted coefficients ($\widehat{B_{ij}}$) and intercepts ($\widehat{\alpha_i}$) for all genes $i$ yields the estimated parameter matrices $\hat{B}$ and $\hat{\alpha}$. All subsequent predictions use these estimated parameters.

#### *In silico* perturbation and prediction

Our objective is to predict the new steady-state transcriptome following a new perturbation that sets the expression of a gene set $I$ to pre-specified values, $x_I$.

First, we partition all genes not in $I$ into two disjoint subsets based on the network topology:

- $U$: the set of genes that are reachable from $I$ via a directed path, where undirected edges are considered traversable in both directions. Their expression levels are altered from the wild-type levels as the perturbation effect propagates through the GRN.
- $K$: the set of genes that are not descendants of any gene in $I$. Their expression is assumed to be unaltered from the wild-type state.

This partitions the full gene set into $U$ (the unknowns to be predicted) and $K^* = K \cup I$ (the knowns). The vector of known expression values $\widehat{x_{K^*}}$ is composed of the pre-specified values for genes in $I$ and the wild-type levels for genes in $K$.

We now use the estimated linear system $X = \hat{\alpha} + \hat{B}X + e$ to predict $x_U$ for perturbed cells. After excluding $x_I$ from the system that is determined by experimental intervention, we have the model in perturbed cells as

$$\begin{pmatrix} \widehat{x_K} \\ x_U \end{pmatrix} = \begin{pmatrix} \widehat{\alpha}_K \\ \widehat{\alpha}_U \end{pmatrix} + \begin{pmatrix} \hat{B}_{KK^*} & \hat{B}_{KU} \\ \hat{B}_{UK^*} & \hat{B}_{UU} \end{pmatrix} \begin{pmatrix} \widehat{x_{K^*}} \\ x_U \end{pmatrix} + \begin{pmatrix} e_K \\ e_U \end{pmatrix}.$$

An observation is $\hat{B}_{KU} = 0$, as any parent of genes in $K$ must reside in $K$, otherwise such a parent in either $I$ or $U$ would create a directed path from $I$ to $K$, contradicting the definition of $K$. Therefore, the only information we have to predict $x_U$ is

$$x_U = \hat{\alpha}_U + \hat{B}_{UK^*}\widehat{x_{K^*}} + \hat{B}_{UU}x_U + e_U.$$

Therefore, we predict $x_U$ by minimizing

$$f(x_U) = \|(I - \hat{B}_{UU})x_U - \hat{\alpha}_U - \hat{B}_{UK^*}\widehat{x_{K^*}}\|^2.$$

If $\hat{B}_{UU}$ has no eigenvalue equal to 1, we have the unique minimizer:

$$\widehat{x_U} = (I - \hat{B}_{UU})^{-1}(\hat{\alpha}_U + \hat{B}_{UK^*}\widehat{x_{K^*}}).$$

#### Interpretation and solution stability

The solution has a clear biological interpretation. The term $\hat{\alpha}_U + \hat{B}_{UK^*}\widehat{x_{K^*}}$ represents the initial, direct regulatory inputs from the known genes $K^*$ onto the unknown genes $U$. The matrix $(I - \hat{B}_{UU})^{-1}$ acts as a propagator, which models the complete propagation of this initial effect through the internal network of the $U$ set. It accounts for all downstream cascades and feedback loops within this subgraph until a new steady state is reached.

The existence of a unique, stable solution depends entirely on the properties of this propagator matrix. A stable equilibrium exists if the spectral radius (the largest magnitude of the eigenvalues) of $\hat{B}_{UU}$ is less than 1 (i.e., $\rho(\hat{B}_{UU}) < 1$). When this condition is met, the propagator matrix can be represented by the Neumann series expansion:

$$(I - \hat{B}_{UU})^{-1} = I + \hat{B}_{UU} + \hat{B}_{UU}^2 + \hat{B}_{UU}^3 + \cdots .$$

This series provides a powerful interpretation: the total effect is the sum of the initial input (multiplied by $I$), plus the effect after one round of propagation through the $U$ network (multiplied by $\hat{B}_{UU}$), plus the effect after a second round ($\hat{B}_{UU}^2$), and so on, until the effect diminishes to zero at a stable equilibrium.

Across all analyses in this study, we have $\rho(\hat{B}_{UU}) < 1$.

### Supplementary Note 4: Simulation details for ATC-PC validation

This note provides the precise data-generating process for the simulation study validating ATC-PC's performance.

We simulated $n = 100{,}000$ cells from two causal topologies where $A \perp\!\!\!\perp C \mid B$: (i) $A \to B \to C$, and (ii) $A \leftarrow B \to C$.

Given a topology, we generated scRNA-seq counts from a Poisson lognormal model. Specifically, under topology $A \to B \to C$, we first simulated latent expression:

$$u_A \sim \mathcal{N}(0, \sigma^2), \quad u_B \sim \mathcal{N}(\alpha_{AB} u_A + s, \sigma^2), \quad u_C \sim \mathcal{N}(\alpha_{BC}(u_B - s), \sigma^2),$$

where $s \in \{0, -2, -4\}$ controls the sparsity level of gene $B$, $\alpha_{AB}, \alpha_{BC}$ were independently drawn from $\mathcal{N}(0, 1)$, and $\sigma = 2$. Then observed counts were simulated as $Y_g \sim \text{Poisson}(\exp(u_g))$ for $g \in \{A, B, C\}$. Similarly, under topology $A \leftarrow B \to C$, the latent expression was simulated as

$$u_B \sim \mathcal{N}(s, \sigma^2), \quad u_A \sim \mathcal{N}(\beta_{BA}(u_B - s), \sigma^2), \quad u_C \sim \mathcal{N}(\beta_{BC}(u_B - s), \sigma^2),$$

where $\beta_{BA}$ and $\beta_{BC}$ were independently drawn from $\mathcal{N}(0, 1)$.

For each topology–sparsity pair, 100 Monte Carlo replicates were generated (600 datasets in total).

### Supplementary Note 5: Construction of gold-standard causal motifs

This note details the procedure for identifying gold-standard causal motifs from single-cell perturbation data, used to validate the ATC-PC method. The same procedure was applied independently to the RPE1 and K562 datasets.

#### Defining high-confidence causal edges

First, we defined a set of high-confidence edges representing strong, direct regulatory effects. An edge from a perturbed gene to a target gene was considered high-confidence if it met all of the following criteria:

- The target gene was significantly differentially expressed upon perturbation of the perturbed gene (FDR-adjusted $p$-value $< 0.001$).
- The absolute Pearson correlation between the expression of the perturbed and target gene was greater than 0.05.
- To focus on the strongest effects, we retained only the top ten target genes for each perturbed gene, ranked by the absolute value of their causal effect size represented by Cohen's D.

#### Constructing causal motifs

Using this set of high-confidence edges, we constructed gene triplets of the below three distinct causal topologies.

- Chain ($A \to B \to C$): We identified triplets with high-confidence edges $A \to B$ and $B \to C$. We enforced that no significant differential expression (DE) signal (FDR-adjusted $p$-value $< 0.05$) existed in any of $B \to A$, $C \to B$, or $C \to A$.
- Fork ($A \leftarrow B \to C$): We identified triplets with high-confidence edges $B \to A$ and $B \to C$. We enforced that no significant DE signal existed in any of $A \to B$, $C \to B$, $A \to C$, or $C \to A$.
- Collider ($A \to B \leftarrow C$): We identified triplets with high-confidence edges $A \to B$ and $C \to B$. We enforced that no significant DE signal existed in any of $B \to A$, $B \to C$, $A \to C$, or $C \to A$.

We further filtered the constructed triplets to ensure their correlation structure matched theoretical expectations:

- For chains and forks, we required the absolute correlation between the non-adjacent gene pair A–C to be weaker than the absolute correlation of both adjacent gene pairs A–B and B–C.
- For colliders, we applied the same filter and additionally required the marginal correlation between $A$ and $C$ to be very weak ($< 0.001$), as expected from their marginal independence.

Finally, to ensure balanced and computationally tractable analysis, if any triplet set contained more than 500 triplets, we randomly sampled it down to 500.

### Supplementary Note 6: Implementation details of GRN inference methods

This note provides the implementation details for all benchmarked GRN inference methods across both the simulation and real-data benchmarks.

**CausalGRN.** We used a significance level of $\alpha = 0.05$ for skeleton inference and $\alpha = 0.05$ for differential expression testing during edge orientation. These parameters were used for both the simulation and real-data benchmarks.

**GES, GIES and PC.** We used the R package pcalg [3] (v2.7.9) for these methods. GES and GIES were run with default parameters, using their respective observational and interventional Gaussian scores. For the computationally intensive PC algorithm, we used a stringent $\alpha = 1 \times 10^{-15}$ in the simulation benchmark, downsampled the wild-type cells for the large K562 and HCT116 datasets to 10,000, and then used $\alpha = 0.01$ for the smaller hESC-DE dataset, $\alpha = 1 \times 10^{-6}$ for the RPE1 and K562 datasets, and $\alpha = 1 \times 10^{-12}$ for the HCT116 dataset due to its stronger gene-gene correlations.

**Lasso-WT and Lasso-All.** We performed Lasso regression using the R package glmnet [4] (v4.1.8). Regularization parameters were selected via five-fold cross-validation.

**GENIE3.** We ran GENIE3 using its R package [5] (v1.24.0) with default parameters. Due to its computational burden, we downsampled the wild-type cells for the large K562 and HCT116 datasets to 10,000 for it.

**GRNBoost2.** We ran GRNBoost2 using the Python package arboteto [6] (v0.1.6) with default parameters.
